# Pesticide contamination of milkweeds across the agricultural, urban, and open spaces of low elevation Northern California

**DOI:** 10.1101/2020.03.09.984187

**Authors:** Christopher A. Halsch, Aimee Code, Sarah M. Hoyle, James A. Fordyce, Nicolas Baert, Matthew L. Forister

## Abstract

Monarch butterflies (*Danaus plexippus*) are in decline in the western United States and are encountering a range of anthropogenic stressors. Pesticides are among the factors that likely contribute to this decline, though the concentrations of these chemicals in non-crop plants is not well documented, especially in complex landscapes with a diversity of crop types and land uses. In this study, we collected 227 milkweed (*Asclepias* spp.) leaf samples from 19 sites representing different land use types across the Central Valley of California. We also sampled plants purchased from two stores that sell to home gardeners. We found 64 pesticides (25 insecticides, 27 fungicides, and 11 herbicides, as well as 1 adjuvant) out of a possible 262 in our screen. Pesticides were detected in every sample, even at sites with little or no pesticide use based on information from landowners. On average, approximately 9 compounds were detected per plant across all sites, with a range of 1 to 25 compounds in any one sample. For the vast majority of pesticides detected, we do not know the biological effects on monarch caterpillars that consume these plants, however we did detect a few compounds for which effects on monarchs have been experimentally investigated. Chlorantraniliprole in particular was identified in 91% of our samples and found to exceed a tested LD_50_ for monarchs in 58 out of 227 samples. Our primary conclusion is the ubiquity of pesticide presence in milkweeds in an early-summer window of time that monarch larvae are likely to be present in the area. Thus, these results are consistent with the hypothesis that pesticide exposure could be a contributing factor to monarch declines in the western United States. This both highlights the need for a greater understanding of the lethal and sublethal effects of these compounds (individually, additively, and synergistically) and suggests the urgent need for strategies that reduce pesticide use and movement on the landscape.

**Contribution to the Field:** Insects are facing multifaceted stressors in the Anthropocene and are in decline in many parts of the world. The widespread use of pesticides is believed to be an important part of the problem. In particular, the monarch butterfly is in sharp decline in the western United States. Here we show that milkweeds in the Central Valley of California, a large urban and agricultural landscape that is part of the monarch breeding and migration route, are contaminated with a diverse array of pesticides. We found a few in high concentrations and many in trace amounts. We do not know how these compounds act together and with other large-scale stressors to cause declines, but it is clear that monarchs and other non-target insects are encountering these pesticides. These results provide critical insight into the growing literature on the impact of pesticides on butterflies specifically and non-target insects more broadly. We hope these field realistic concentrations will aid in the design of further experiments in the field and the lab.

## Introduction

Widespread reports of declining insect populations have received considerable and increasing attention in recent years (Forister et al., 2010; Potts et al., 2010; Hallmann et al., 2017; Janzen and Hallwachs, 2019; Sánchez-Bayo and Wyckhuys, 2019; Wepprich et al., 2019). The causes of this phenomenon are multi-faceted, as species face correlated anthropogenic stressors that include climate change, habitat loss, and the use of pesticides (Deutsch et al., 2008; Goulson et al., 2015; Forister et al., 2019; Sánchez-Bayo and Wyckhuys, 2019). While the importance of each of these drivers will vary with context, just one or a combination of factors can disrupt population dynamics and lead to extirpation or extinction (Brook et al., 2008; Tylianakis et al., 2008; Potts et al., 2010; González-Varo et al., 2013). One potentially devastating combination of stressors is the historical loss of habitat to agricultural intensification and the contemporary use of pesticides on modified lands (Gibbs et al., 2009). To better understand the contribution of pesticides to long-term trends in insect populations, especially in heavily converted landscapes, we must identify the diversity of compounds, quantify their concentrations, and test how these affect insect survival and performance. Here we investigate the suite of pesticides that potentially contaminate milkweeds in the Central Valley of California, a large agricultural and urban landscape. It is our intention that the results reported here will provide critical data on field-realistic concentrations of pesticides in modified landscapes in order to better parameterize laboratory experiments on pesticide toxicity affecting non-target organisms.

Pesticides have long been discussed as drivers of ecosystem disruption and insect declines, especially in the context of agriculture (Epstein, 2014). Conventional agriculture employs a wide range of pesticides (including herbicides, insecticides, and fungicides) which can affect both target and non-target species (Pisa et al., 2014; Abbes et al., 2015). Insecticides and fungicides can have direct effects on insects (Sanchez-Bayo and Goka, 2014; Mulé et al., 2017), while herbicides are most often associated with indirect effects by altering the nearby plant community and floral resources, though some recent research indicates certain herbicides can also have direct effects on insects (Egan et al., 2014; Balbuena et al., 2015; Dai et al., 2018; Motta et al., 2018). Recently much attention has been paid to neonicotinoids, a class of anticholinergic insecticides, whose use has dramatically increased over the past 20 years, such that they are now the most widely used class of insecticide in the world (Wood and Goulson, 2017). Neonicotinoids are water-soluble and readily taken up by plant tissues, posing a risk to non-target insects as they can be found in all plant parts, including leaves, pollen and nectar (Bonmatin et al., 2015; Wood and Goulson, 2017). Much research has focused on their impacts on bees (Whitehorn et al., 2012), however their use is also associated with declines of butterflies in Europe (Gilburn et al., 2015) and in the Central Valley (Forister et al., 2016). While individual pesticides can have lethal and sub-lethal effects (Pisa et al., 2014), plants sampled in agricultural landscapes often contain multiple compounds (Krupke et al., 2012; Olaya-Arenas and Kaplan, 2019). The literature on the additive or synergistic effects of pesticide combinations on non-target organisms is sparse, however particular combinations have been shown to behave synergistically in insects broadly (Zhu et al., 2014; Morrissey et al., 2015) and pest Lepidoptera specifically (Jones et al., 2012a; Liu et al., 2018a; Chen et al., 2019). By focusing on one or a few select pesticides or even a single class of pesticides, the realized risk of these chemicals on non-target insects is likely being underestimated.

Perhaps the most noted recent decline of any insect is that of the monarch butterfly (*Danaus plexippus*), whose reduced numbers have been observed in both the eastern (Stenoien et al., 2018) and western (Espeset et al., 2016; Schultz et al., 2017) North American populations. In the eastern United States, many hypotheses have been proposed to explain the monarch decline, including loss of critical overwintering habitat, natural enemies, climate, and various pesticides, especially herbicides, that have reduced milkweed abundance (*Asclepias* spp.) (Belsky and Joshi, 2018). In the west, monarch overwintering populations reached a historic low in 2018 (Pelton et al., 2019), and the causes appear to include loss of overwintering habitat, climate, and pesticides (Crone et al., 2019). There are few studies evaluating the direct (lethal and sub-lethal) effects of pesticides on the monarch (Krischik et al., 2015; Pecenka and Lundgren, 2015; James, 2019; Krishnan et al., 2020). Pecenka and Lundgren tested the toxicity of the neonicotinoid clothianidin and observed it in sub-lethal concentrations in milkweeds sampled in South Dakota, U.S.A (Pecenka and Lundgren, 2015). Krischik et al. (2015) and James (2019) both assessed the effects of imidacloprid on monarchs. Krishnan et al. (2020) investigated the toxicity of five compounds on larval monarchs, including chlorantraniliprole, imidacloprid, and thiamethoxam. Further work in the mid-western U.S. sampled milkweeds and screened leaf samples for pesticides (Olaya-Arenas and Kaplan, 2019). A total of 14 pesticides were identified at various concentrations, including clothianidin, which was found in similar concentrations as those reported by Pecenka and Lundgren (2015). While these findings show that pesticides can be found at physiologically relevant concentrations in milkweeds in the eastern United States, we currently lack an understanding of pesticide contamination in the west and thus have no direct way to assess the potential contribution of pesticides to the decline of the western monarch.

The Central Valley of California is the largest cropped agricultural landscape of the western United States and is part of the migratory distribution and breeding ground for the western population of the monarch butterfly. Historically, one of the primary anthropogenic stressors in the Central Valley has been the loss of wetland habitat to agricultural intensification (Reiter et al., 2015). This change to the landscape reduced floral resources and introduced pesticides to large portions of the landscape (Wagner, 2019). While a major contributor, agriculture is not the only source of pesticides in the environment as pesticides are commonly sold for home and garden use (Atwood and Paisley-Jones, 2017). Over the past three decades the Sacramento Valley, the largest metropolitan area in the Central Valley, has become increasingly developed (Theobald, 2005) and this urban growth may represent a second major source of contaminants in the region (Weston et al., 2009). Considering the history of the region, monarchs and other native and beneficial insects may be encountering a heterogeneous and toxic chemical landscape.

In this study, we measured the concentration and diversity of pesticides found in *Asclepias* spp. leaves collected in the Central Valley of California. Over four days in late June of 2019, we sampled leaves from different land use types, including agriculture, wildlife refuges, urban parks and gardens, and plants sold in retail nurseries. The first objective of this study is to gather a snapshot picture of which pesticides are present on the landscape and in what concentrations they are found when monarch larvae are expected to be feeding. Second, we present an exploratory examination of contamination differences among land use types. Finally, we ask if the contamination levels detected could harm monarchs or other terrestrial insects, based on published data. Thus, this study is designed as a first look into what pesticides monarch larvae might be exposed to in the Central Valley and not to directly test if they are responsible for the ongoing decline of the western population.

## Methods

### Milkweed sampling

Milkweed samples of *Asclepias fasicularis* (161 samples) and *A. speciosa* (50), with fewer *A. eriocarpa* (4) and *A. curassavica* (12), were collected from sites in the Central Valley and purchased from retail nurseries from June 24-27, 2019 (fig. 1A). Our collection time was intended to overlap with monarch breeding in the Central Valley based on personal observations and historical data (Espeset et al., 2016). In total we collected samples from 19 different sites: five sites were located in conventional farms, one in an organic farm, one in a milkweed establishment trial (grown for restoration), one in a roadside location (adjacent to an agriculture field), five in wildlife refuges, four in urban areas, and two from retail nurseries. The agricultural locations (including the restoration trial and the roadside location) were all treated in analyses as “agriculture” (since replication was not sufficient to parse further); thus our main land type categories were “agriculture”, “refuge”, “retail” and “urban.” Sites were selected opportunistically, based on accessibility and in order to sample a diversity of landscapes. The identity of the milkweed species is mostly confounded with sampling location (Table S1), so our inferential ability is limited for differences in contamination among plant species. If sites contained fewer than 20 plants, all plants were surveyed and if sites contained greater than 20 plants, individual plants were selected randomly within each patch, and leaf samples were collected and placed in bags. Clippers were cleaned with rubbing alcohol between every cutting. Samples were transported on ice, frozen and stored, and ultimately shipped to the Cornell University Chemical Ecology Core Facility lab on dry ice.

**Figure 1.**
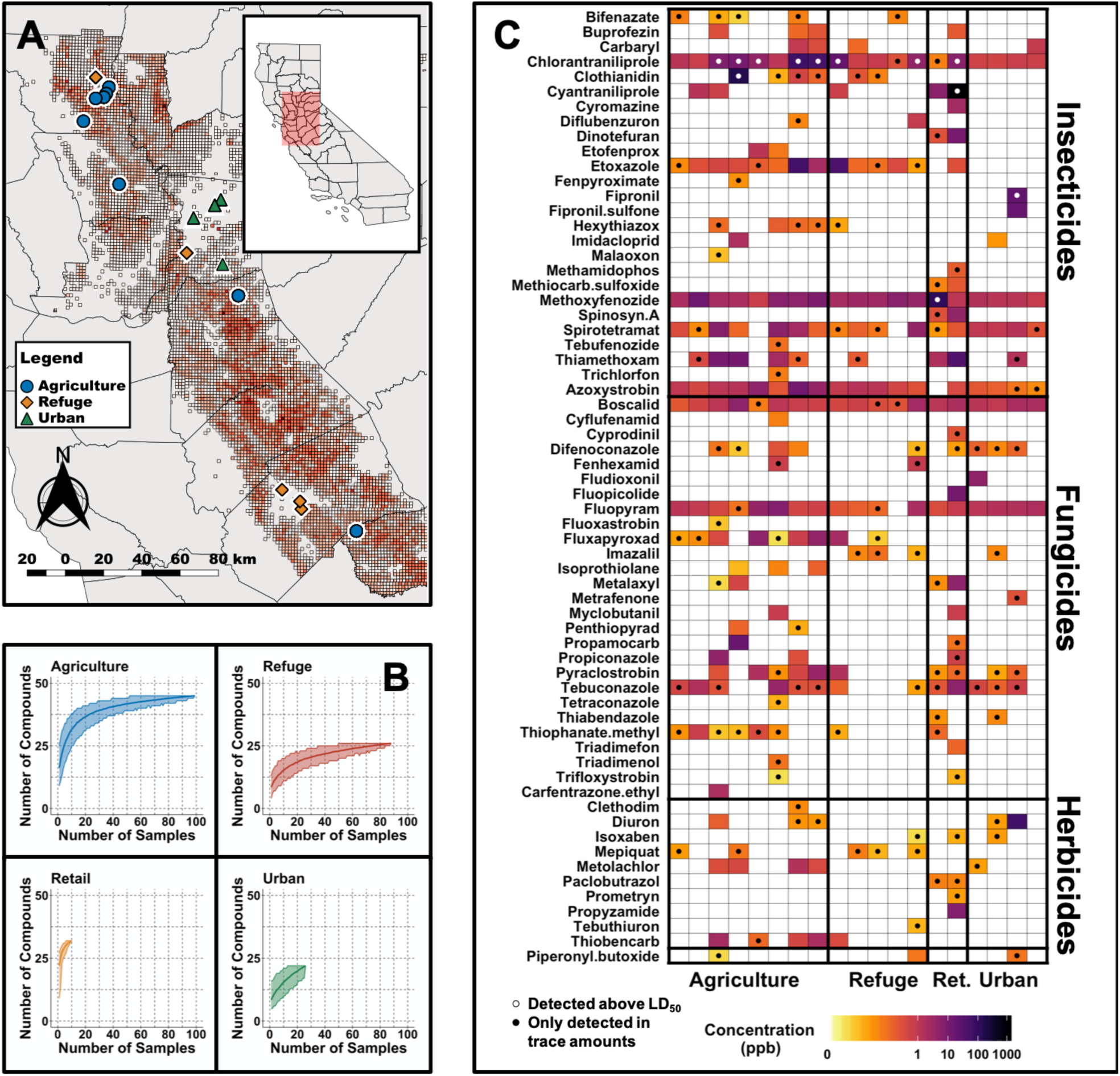
Overview of pesticide compounds detected in the Central Valley. A) Sampling locations colored by land use type. Red background indicates the number of compounds reported in the 2015-2017 California Department of Pesticide Regulation pesticide use data (the range is from 1 compound for the lightest gray to 113 for the darkest red cells). B) Rarefaction curves for the number of pesticides detected by land use type. C) Mean concentrations (per plant) of compounds at each site. Values are shown in parts per billion on a log scale. Black circles indicate compounds only detected in trace amounts (i.e. below the level of quantification). White circles indicate compounds found above a lepidopteran LD_50_.

### Chemistry

Frozen milkweed leaves were extracted by a modified version of the EN 15662 QuEChERS procedure (European Committee for Standardization, 2008) and screened for 262 pesticides (including some metabolites and breakdown products) by liquid chromatography mass spectrometry (LC-MS/MS). Five grams of frozen leaves (5 grams was the target sample weight, samples ranged from 0.35 to 5.07 grams and were prepared accordingly) were mixed with 7 mL of acetonitrile and 5 mL of water. The leaves were then homogenized for 1 min using ceramic beads (2.8 mm diameter) and a Bead Ruptor 24 (OMNI International, USA). After complete homogenization, 6.5 mg of EN 15662 salts were added (4 g MgSO4; 1 g NaCl; 1 g sodium citrate tribasic dihydrate; 0.5 g sodium citrate dibasic sesquihydrate). Samples were then shaken and centrifuged at 7300 × g for 5 minutes. One milliliter of supernatant was collected and transferred into a d-SPE (dispersive solid phase extraction) tube containing 150 mg PSA, 900 mg MgSO4. After the d-SPE step, 496 µL of supernatant were collected and 4 µL of a solution of 5 internal standards spanning across a wide range of polarity (d4-imidacloprid 0.07 ng/µL; d10-chlorpyrifos 0.2 ng/µL: d7-bentazon 0.1 ng/µL; d5-atrazine 0.02 ng/µL; d7-propamocarb 0.1 ng/µL) was added. Samples were then filtered (0.22 µm, PTFE) and stored at −20°C before analysis.

Sample analysis was carried out with a Vanquish Flex UHPLC system (Dionex Softron GmbH, Germering, Germany) coupled with a TSQ Quantis mass spectrometer (Thermo Scientific, San Jose, CA). The UHPLC was equipped with an Accurcore aQ column (100 mm × 2.1 mm, 2.6 µm particle size). The mobile phase consisted of (A) Methanol/Water (2:98, v/v) with 5 mM ammonium formate and 0.1% formic acid and (B) Methanol/Water (98:2, v/v) with 5 mM ammonium formate and 0.1% formic acid. The temperature of the column was maintained at 25°C throughout the run and the flow rate was 300 µL/min. The elution program was the following: 1.5 min equilibration (0% B) prior to injection, 0-0.5 min (0% B, isocratic), 0.5-7 min (0%-70% B, linear gradient), 7-9 min (70%100% B, linear gradient), 9-12 min (100% B, column wash), 12-12.1 min (100%-0% B, linear gradient), 12.1-14.5 min (0% B, re-equilibration). The flow from the LC was directed to the mass spectrometer through a Heated Electrospray probe (H-ESI). The settings of the H-ESI were: spray voltage 3700 V for positive mode and 2500 V for negative mode, Sheath gas 35 (arbitrary unit), Auxiliary gas 8 (arbitrary unit), Sweep gas 1 (arbitrary unit), Ion transfer tube temperature 325°C, Vaporizer temperature 350°C. The MS/MS detection was carried out using the Selected Reaction Monitoring (SRM) mode. Two transitions were monitored for each compound: one for quantification and the other for confirmation. The SRM parameters for each individual pesticide are summarized in Table S2. The resolution of both Q1 and Q3 was set at 0.7 FWHM, the cycle time was 0.5 s and the pressure of the collision gas (argon) was set at 2 mTorr.

### Statistical analyses

The chemical screening was able to classify concentrations into four categories. The first was when the chemical was below the level of detection and these were treated as zeros. Second was when the chemical was detected, but the concentrations were low to be quantified, these samples were labeled as “trace”. In these cases, we used a known lower limit of detection for the observed value. Third was if the chemical could be detected and quantified. Finally, there were a few cases in which chemicals were found in too high of concentrations to be quantified. In these cases, we used the upper limit of detection as the observed value. The lower and upper limits of detection are known values which vary by compound, thus even if a compound was only found in trace amounts, we can still draw some inference about relative concentrations.

Sampling sites were classified into agricultural, retail, refuge, or urban for statistical analysis, as described above. To examine total pesticide richness and diversity in each land use type we performed sample-based rarefaction. To directly compare compositional differences in pesticides between different land use types, we calculated the effective number of pesticides for each sample using different Hill numbers (*q* = 0, *q* = 1, and *q* = 2). Using this approach to diversity, the sensitivity to rare compounds changes as a function of the parameter *q*: *q* = 0 weights all compounds equally (richness), *q* = 1 weights all compounds by their relative abundance (exponential of Shannon entropy), and q = 2 down-weights rarer compounds (inverse Simpson’s index) (Hill, 1973; Jost, 2006). We also performed this same diversity analysis, but on data that were rarefied to match the land use type with the lowest sampling effort (retail, 11 samples).

Dissimilarity of pesticides detected among milkweeds from each of the habitat types was then visualized using a distance-based redundancy analysis (dbRDA) (Legendre and Legendre, 2012). The distance matrix was constructed using the quantitative generalization of Jaccard dissimilarity (Ružička index) with habitat types as the constraining factors (Schubert and Telcs, 2014). The dbRDA was implemented using the R package Vegan (Oksanen et al., 2019). Associations between each pesticide and habitat type were examined using the group-equalized point serial correlation (De Cáceres and Legendre, 2009). We explored associations allowing pesticides to be indicative of combinations of habitat types (De Caceres et al. 2010). Statistical significance (*α* = 0.05) of the strongest association for each pesticide with land type was determined using 9999 permutations of the data. These indicator analyses were conducted using functions from the R package indicspecies (De Cáceres et al., 2020).

### Literature search

To examine biological importance of the detected concentrations, we compared our findings to published LD_50_ data for honeybees and Lepidoptera. LD_50_ data (both contact and oral where available) for honeybees were collected from EPA records in the ECOTOX and Pubchem databases and the University of Hertfordshire’s Pesticide Properties Database (Table S3). One strength of these data is their standardized collection and thus ease of use for comparison across compounds in examining collective (or additive) effects. To do this, we calculated the hazard quotient for each compound, by dividing the detected concentration by the LD_50_, and then summed this across all compounds in each sample (Stoner et al., 2019). This approach has an important drawback in that assumes a linear relationship between concentration and effect, which is often not realized, and then propagates this across all compounds in each sample. Compounds may have no effect or a different effect at trace concentrations compared to a LD_50_; however, this calculation assumes they have an effect proportional to the LD_50_. Additionally, while the EPA uses honeybees as a surrogate species for all non-target terrestrial invertebrates in pesticide risk assessments, these data are not directly applicable to lepidopterans and many other insects. Furthermore, toxicity tests are performed on adult honeybees which are of course different from larval Lepidoptera, and this is especially true considering that some insecticides are designed specifically to affect juvenile lepidopterans. We only use the honeybee LD_50_ data in the most general sense to establish a benchmark of the concentrations where these compounds can have a biological effect on non-target terrestrial invertebrates. To better apply our findings directly to the monarch butterfly, we also conducted a literature review of papers that have studied the compounds we detected and have reported LD_50_ concentrations for lepidopterans (Table S4). The literature search was performed in January 2020 using ISI Web of Science with the terms (lepidopt* OR butterfly* OR moth*) and (compound) and was repeated for all compounds identified in our samples.

## Results

A total of 64 compounds were identified in at least one leaf sample out of 262 possible compounds in our test panel. Of these, 25 were insecticides (including two insecticide metabolites), 27 were fungicides, 11 were herbicides, and 1 was a common adjuvant (fig. 1C). An adjuvant is a compound designed to enhance the effect of other compounds. Seven compounds were detected in over 50% of collected samples and seventeen compounds were detected in over 10% of samples. Methoxyfenozide and chlorantraniliprole were the most prevalent compounds, which were found in 96% and 91% of samples respectively. Detected concentrations across all compounds range from below 1 ppb to above 900 ppb. In some samples, compounds were detected, but the concentration was too low to be quantified (fig. 1C). In these cases, we used the limit of detection value for that pesticide, as the actual concentration would be above the limit of detection but below the limit of quantification.

Generally, more pesticides were found in agricultural and retail samples than refuge or urban samples, however we detected considerable variation and pesticides were present in all land use types (fig. 1B, fig. 2, fig. S1). Diversity analyses show especially high numbers of compounds in retail samples, and this appears to be driven by “rare” compounds (found in only one or a few samples), as effective numbers of compounds dramatically decline between Hill numbers generated at *q*= 0 and q=1 (fig. 2). The other three land use types contain a similar proportion of common to rare compounds. This pattern is maintained even when samples are rarefied to match the low sampling effort of the retail samples (fig. S1). There was substantial variation in the mean number of compounds among milkweed species, however, as previously noted, species are confounded with sampling sites as most sites had only one species present (fig. S2). This is especially true for *Asclepias curassavica* and *Asclepias eriocarpa*, which were almost exclusively found in retail and agricultural sites respectively (Table S1). When examining site dissimilarity across all compounds, there is clustering based on land use type in ordination space (Fig. 3). In general, retail and agricultural samples are the most similar, but there are also refuge sites that are chemically similar to agriculture and retail sites (fig. 3). Many specific chemicals are associated with agricultural sites including chlorantraniliprole, clothianidin, imidacloprid, and azoxystrobin (fig. S3, Table 1). Methoxyfenozide and thiamethoxam are associated with retail samples, however it is important to note the low sample size of retail compared to other land use types. We have stronger evidence supporting associations with agriculture than associations with retail.

**Table 1.**
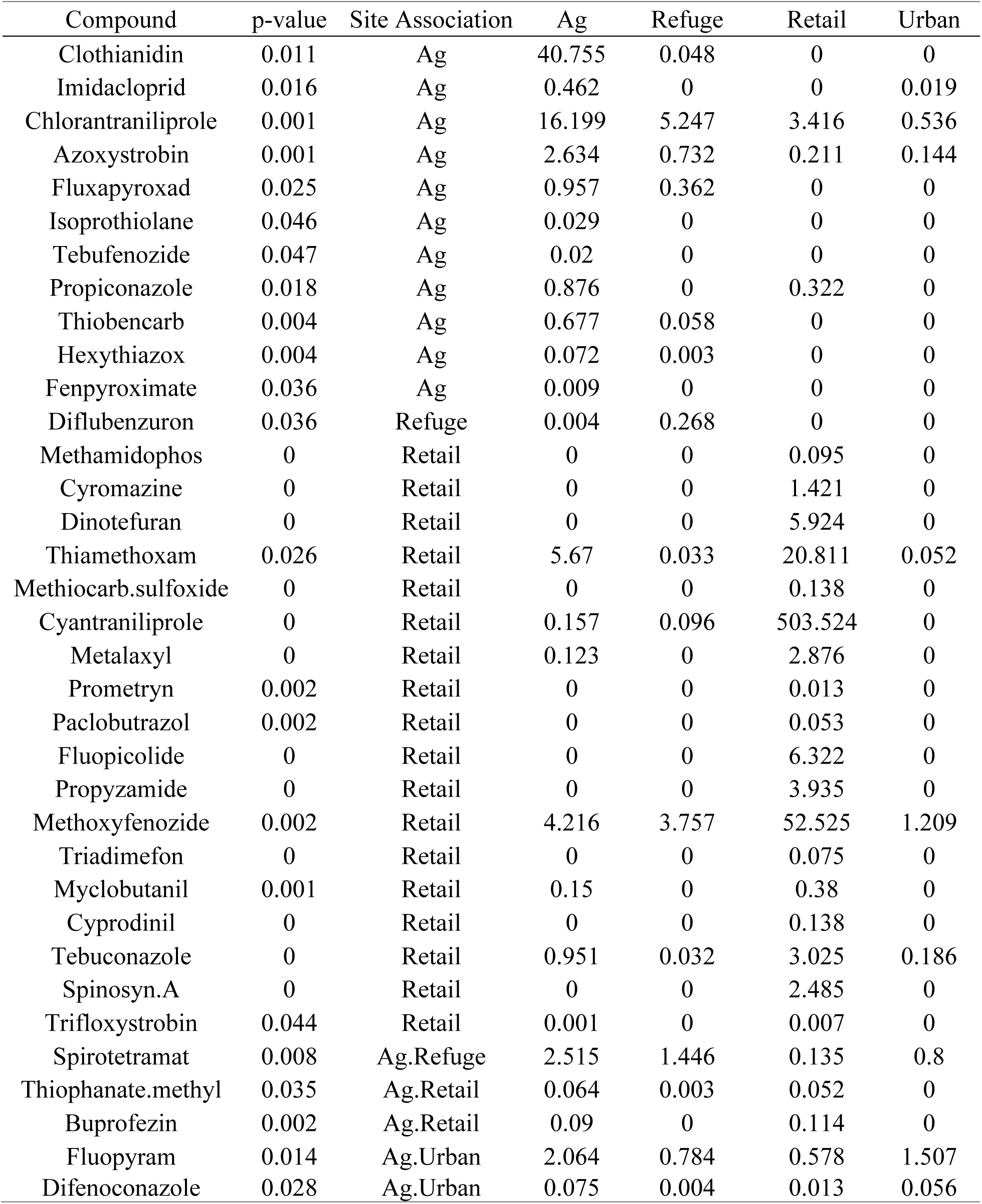
Indicator species results for associations between sites and individual compounds. Values in each land type category show mean concentration (ppb). Only “significant” relationships (at α = 0.05) are shown. No corrections were made for multiple comparisons.

**Figure 2.**
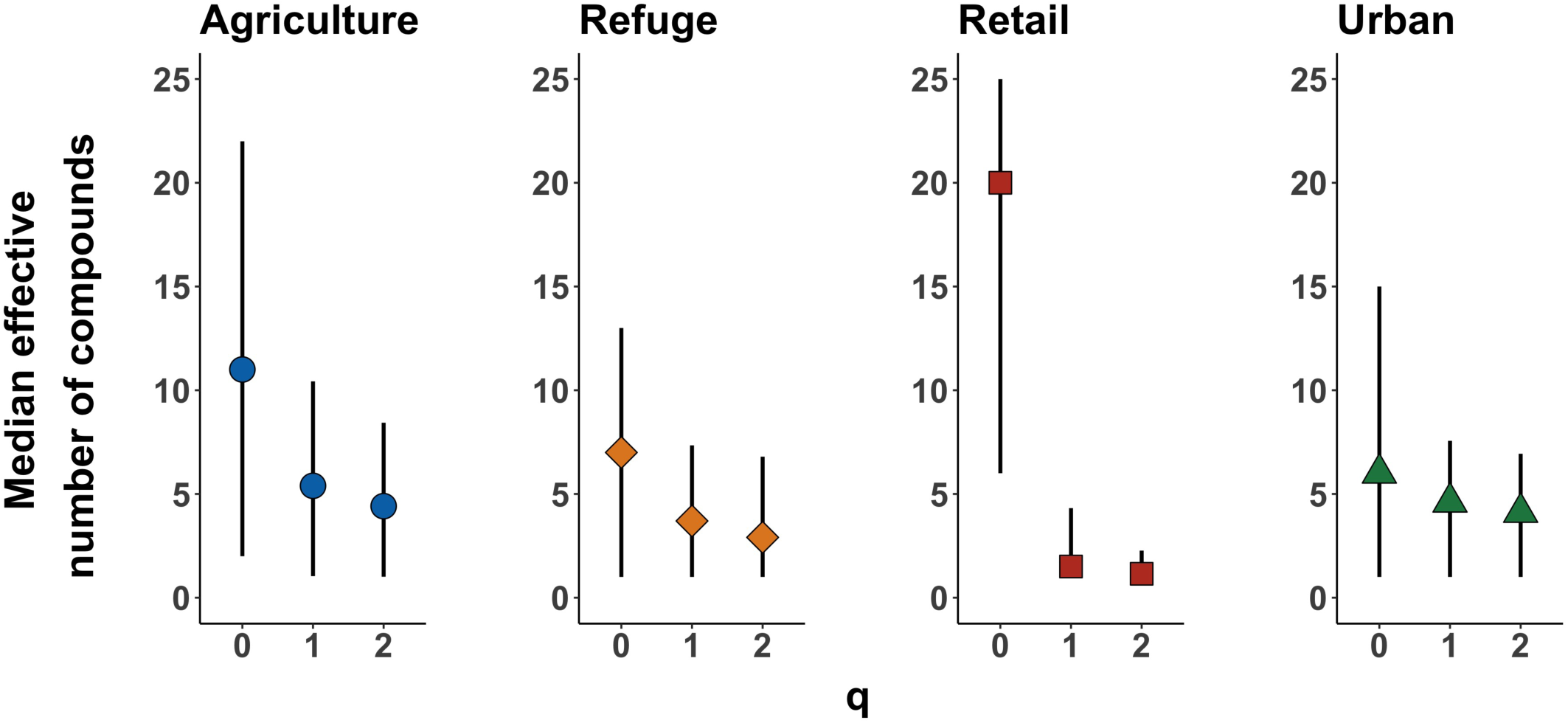
Effective numbers of pesticides per sample by land use type using Hill numbers generated across a range of *q* values that place different weights on rare vs common compounds (at *q* = 0 all compounds have equal weight, see text for additional details). Points show the median number of compounds per sample. Bars show the full range across samples within one land use type.

**Figure 3.**
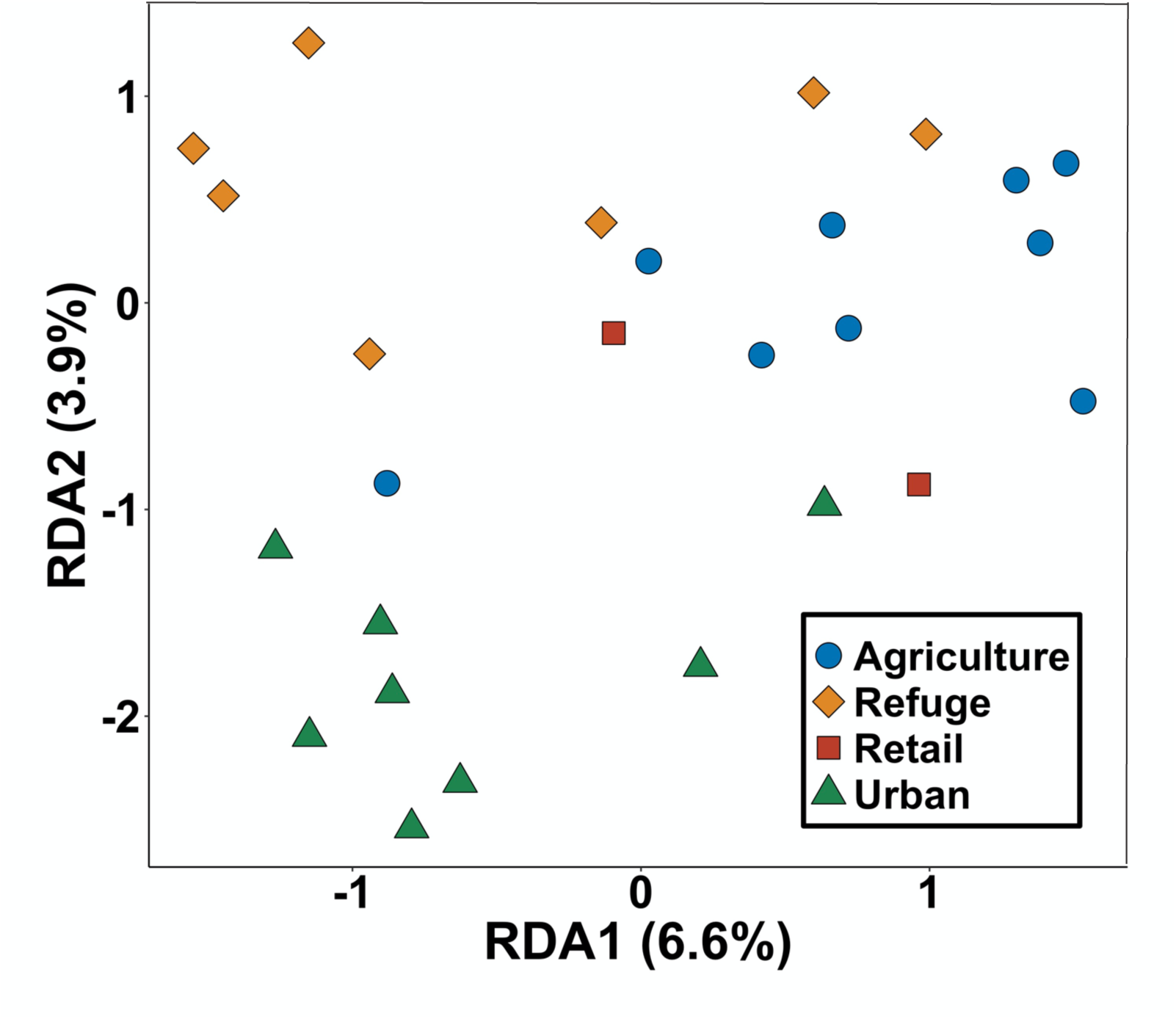
Ordination of the constrained axes from distance-based redundancy analysis based upon chemical dissimilarity between sampling sites (variation explained by axis indicated after each axis label). Points indicate the mean score for each sampling site; colors and shapes indicate land use type.

Of the 64 detected compounds, we acquired contact and oral honeybee LD_50_ concentrations for 62 compounds (data were not available for the two insecticide metabolites). When looking at each compound individually, there were 27 exceedances of a contact LD_50_ and 52 exceedances of an oral LD_50_. These 79 total exceedances occurred in 36 individual plant samples (out of 227) from seven sites. Calculating collective risk across all detected compounds in a sample (by dividing the observed value by the LD_50_ and then summing across the sample) identified the same 36 samples, thus it appears individual compounds are driving the exceedances of honeybee LD_50_ concentrations. These samples primarily came from agricultural or retail samples, however one urban backyard sample also exceeded an oral LD_50_. Information about exceedances of specific compounds can be found in a supplemental table (Table S5).

The literature search for Lepidoptera and pesticides generated 44 studies with published lethal doses for the compounds we detected (Table S4). Pest species dominated the literature as only 8 non-pest papers (including the 4 aforementioned monarch papers) were found. The majority of compounds had none or a single study. Reported LD_50_ concentrations for a compound often varied between lepidopteran species by multiple orders of magnitude. Generally, insecticides had lower LD_50_ values (and thus are more directly toxic) than fungicides and herbicides. An additional axis of variation in the literature was exposure time, which varied from under 30 minutes to three weeks, however the range of 24-72 hours was most common. Using the published lepidopteran data, 47% of samples exceeded published LD_50_ values for a lepidopteran. Of these, 68% (32% of all samples) contained a pesticide above a published LD_50_ for monarchs. These exceedances were observed in 10 sites across all land use types, however agriculture and refuge contained the highest number of raw exceedances (they are also the most sampled) (fig 1B). The most notable individual compound is chlorantraniliprole, which was found above a published LD_50_ for monarchs in 26% of all samples and above an LD_10_ in 78% of all samples. Clothianidin was recorded above a monarch LD_50_ in 15 samples (and above the LD_90_ in 11), however these all came from one agricultural site. Other compounds that exceeded an LD_50_ were cyantraniliprole, fipronil, and methoxyfenozide which came from retail and urban samples. A full overview of all of the exceedances and their associated land use type can be seen in figure 1.

## Discussion

Insects are facing many stressors simultaneously, especially in areas where habitat has already been converted from a natural state and fragmented. Identifying various stressors and quantifying their implications for population dynamics are critical for fully understanding how insects are responding to the Anthropocene. In the Central Valley, pesticides likely represent an important stressor, as they were detected in all land use types sampled. Agricultural and retail samples tended to have more compounds in higher concentrations, however our choice of sampling locations was not random, nor comprehensive, and thus our ability to make direct land use type comparisons is limited. In general, we suspect that our results may be conservative. Agricultural samples were primarily collected from farmers who are already working with the Xerces Society to implement on-farm invertebrate conservation, many of whom have made an effort to avoid bee-toxic pesticides. Likewise, the backyard samples were taken at the homes of Xerces employees where pesticides have not knowingly been applied recently. Still, both of our backyard sites had pesticide detections, including one site with residues from an application of fipronil made more than six years before sampling. Numerous pesticides were also detected in wildlife refuges, though some herbicides known to be used on portions of the refuges were not detected. All of the refuges sampled are surrounded by agricultural fields. In combination with the backyard samples, this demonstrates the presence of pesticides in areas where they are not expected or generally used and are likely coming from adjacent areas.

Another reason to suspect that our results are conservative comes from the chemical screening process itself. There are several pesticides that would likely have been identified if they had been part of the panel that was used in screening. Pyrethroid insecticides, including bifenthrin, and some fungicides, including chlorothalonil, could not be detected with the lab methods used, but are commonly applied to crops in the Central Valley and are toxic to non-target insects (Wolfenbarger et al., 2008). Overall, the clearest pattern in these data is the ubiquity of pesticide presence in milkweeds across the Central Valley, which may impact local and migratory insects (monarch caterpillars are not the only insects that interact with these plants) as they are very likely being exposed to many contaminants. This is true whether a caterpillar is consuming a milkweed leaf in a wildlife refuge, a backyard, or near a conventional agricultural field.

While compounds and concentrations were highly variable, a few notable pesticides warrant further discussion. Chlorantraniliprole was the second most common pesticide, identified in 91% of samples. Krishnan et al. recently studied the toxicity of this specific compound in different instars of monarchs (Krishnan et al., 2020). They found chlorantraniliprole to be highly toxic when compared to imidacloprid and thiamethoxam. Chlorantraniliprole’s LD_50_ was lowest (and thus most toxic) in second instar caterpillars. The number of exceedances we report for this compound used this second instar value. We also found a high number of exceedances of the reported LD_10_ for second instars. These lower doses are often used as a benchmark for sub-lethal effects (Perveen, 2000; Hummelbrunner and Isman, 2001), thus raising the possibility that the majority of our samples contained residues of chlorantraniliprole that could impact the biology of the overall monarch population, while not directly causing mortality. Clothianidin was detected well above lethal concentrations for larval monarchs at one site. It is interesting to note that we have anecdotally linked this finding to an application in the weeks preceding sampling by the landowner to a nearby field, thus providing further evidence of movement of compounds on the landscape. Another compound of note was methoxyfenozide, which was the most frequently detected compound across samples. This compound is an insect growth regulator that targets juvenile lepidopteran pests and is commonly applied during May, June and July in the counties we sampled. Methoxyfenozide accelerates molting in lepidopteran species, and while they have not been directly tested, monarch butterflies have been predicted to be susceptible to this class of pesticides (LaLone et al., 2014). Bees are not predicted to be as sensitive to methoxyfenozide, suggesting that using honeybee data as a surrogate would underestimate risk to non-target juvenile lepidopterans such as monarchs.

There are some notable caveats when applying the above studies to our findings. First, these studies exposed caterpillars at various instars and for different exposure times. It is not clear how an LD_50_ of one compound over 36 hours compares with an LD_50_ of a different compound over 48 hours, and what either of these can tell us about risk in the field. A larval monarch will consume a plant for much longer than 48 hours, and generally longer exposure times will decrease survival (Abivardi et al., 1999; Yue et al., 2003; Wang et al., 2009, 2013; Nasr et al., 2010; Rehan and Freed, 2015; Ahmed et al., 2016; Liu et al., 2018b). Thus, considering shorter exposure times is likely to be a conservative approach which underestimates risk. We should also consider temporal issues from the perspective of plants. Pesticides are not static in leaves and concentrations will dissipate over time. The half-lives for some of these compounds have been investigated in different plants and there is high variation (Fantke and Juraske, 2013; Fantke et al., 2014). Reported half-lives range from shorter than a day to longer than the life of a monarch caterpillar. Given that the LD_50_ values we obtained have shorter exposure lengths compared to the feeding time of monarch caterpillars, these LD_50_ values may better account for reduced exposure due to pesticide turnover in plant tissue. Additionally, our sampling timing certainly impacted the chemicals and concentrations we found. It is likely that we would have detected different pesticides had we sampled in late July or August instead of June. It is important to note that we specifically planned our timing to be during the period that a larval monarch could be present in the Central Valley. A final point of uncertainty worth noting is behavior: monarchs are known to express oviposition preferences among different species of milkweed (Pocius et al., 2018), but it is currently unknown whether pesticide contamination can be a factor in this decision. Despite these uncertainties, we think that these reported LD_50_ concentrations offer compelling evidence that certain compounds are being found at biologically meaningful concentrations, with possible regicidal (or sub-regicidal) implications for larval monarchs in the Central Valley.

With the exception of the already mentioned compounds, we are not able to speculate how the concentrations we detected for most compounds directly impact larval monarchs. Overall, most of the concentrations we observed were below reported LD_50_ values for other lepidopterans and honeybees, however there are numerous reasons why most reported LD_50_ values may not be reliable for monarchs or other non-target butterflies and moths. The vast majority of studies on the compounds we detected are focused on lepidopteran pest species, and many of these studies investigate lethal concentrations on populations suspected to display pesticide resistance. A study on a resistant population will inflate the reported lethal doses and, thus, these studies likely do not reflect the risk of pesticides for non-target insects. Additionally, most studies have the same exposure time drawback already discussed, namely short exposures. This common experimental design is ideal for determining how to deter pests with a minimal number of applications, however it is not a good benchmark for understanding how lethal these contaminants are to non-target insects. It is critical that future research continues to quantify toxicity of these compounds, for monarchs and other insects for which we currently have no data.

Moving beyond individual compounds, these findings raise the possibility of harmful effects from combinations of multiple compounds, even if each is present at low levels. We explored collective (or additive) effects of compounds using honeybee data, which are highly standardized and allow for comparison of compounds within one sample. High risk samples were typically driven by a single compound in high concentration with little contribution from all of the others. We have already stated the assumed linear relationship of this calculation and the lack of applicability of bee data for larval caterpillars, but this allowed for some quantification of collective effects. This does not mean that the low concentration of many compounds is not important, as they could act synergistically, which cannot be quantified with the current data. There are far fewer studies on interactions of multiple compounds, however synergistic effects have been identified in Lepidoptera for thiamethoxam, chlorantraniliprole, imidacloprid, and methoxyfenozide (Jones et al., 2012b; Liu et al., 2018b; Chen et al., 2019), all of which we detected. These findings suggest possible negative effects on lepidopterans; however, it is clear that more research is needed to understand the synergistic effects of field-relevant concentrations on non-target insects.

This is now the second study in the past year that has found pesticide contamination in milkweeds that could be used by breeding monarchs. Olaya-Arenas and Kaplan (2019) also found that pesticides were present in milkweed samples collected near agricultural fields in the mid-western U.S. That study found a total of 14 compounds, however the authors screened for different and fewer compounds than this study. When directly comparing 30 compounds that both studies looked for, Olaya-Arenas and Kaplan found 12 compounds while we detected 14 out of 30. This result is unexpected as that study was concentrated in corn and soybean fields, while our study covered many different land use types and agricultural areas with higher crop diversity. That study collected more than five times as many samples over two years, which may account for the similar number of compounds despite less land use diversity. Similar to our study, Olaya-Arenas and Kaplan were not able to definitively conclude that the pesticides they observed are negatively impacting monarchs, as we currently lack the appropriate data, however it is likely that they are encountering biologically meaningful concentrations of these contaminants in the landscape.

Pesticides are frequently discussed as a driver of insect declines, which have been reported in the Central Valley for butterflies in general (Forister et al., 2010) and for monarchs in particular (Espeset et al., 2016). Notably, while monarchs are in decline in the region, many other butterfly species show even steeper declines (Nice et al., 2019). We are not suggesting that pesticides are solely responsible or even the most important factor in these declines, however our findings demonstrate the potential for pesticides to play a role. Insecticides, fungicides, and herbicides were found in milkweeds at all sampling sites, even in locations we know have not been directly treated. Compounds were also detected in milkweeds purchased from commercial suppliers used by the general public for plantings intended to support butterfly conservation. We are not aware if our findings apply to other butterfly host plants in the region, however given our knowledge that many of these exposures are caused by off-site movement, similar contamination can be expected on other plants found throughout this highly developed landscape. Much more research will be needed to understand how these different concentrations impact monarchs (and other pollinators and beneficial insects) and we hope that our data provide a useful starting place for future experimental designs. We also hope that the results presented here emphasize the need for additional research on practices that reduce pesticide use and movement across landscapes with many uses, including habitat for native insects.

## Acknowledgements

We thank Linda S. Raynolds for generous support of this project, Maggie Douglas for help in gathering the bee toxicity data, and Ian Kaplan, Tom Dilts, and Jaret Daniels for discussion of results. M.L.F was supported by a Trevor James McMinn professorship.

## Contribution of authors

C.A.H., S.M.H., and other Xerces staff collected the samples. N.B. performed the chemical analysis. J.A.F., C.A.H., and M.L.F performed statistical analyses. All authors wrote the manuscript.

## Supporting information

Table S1. Number of samples of different milkweed species from different land use types.

Table S2. Retention times and optimized SRM acquisition parameters for pesticides and internal standards (RT: Retention time, CE: Collision Energy)

Table S3. Contact and oral LD_50_ data for honeybees.

Table S4. Studies from Lepidoptera literature review.

Table S5. Exceedances of honeybee LD_50_ concentrations by land use type and compound.

Table S6. Indicator species results for associations between sites and individual compounds. Values in each land type category show mean concentration (ppb).

Figure S1. Mean effective numbers of pesticides per sample by land use type using different hill numbers after rarefaction. Points show the mean effective number of compounds per sample. Error bars show the range of effective numbers of pesticides across samples within one land use type.

Figure S2. Variation in the number of compounds per sample by milkweed species. Bars show the maximum and minimum number of compounds detected in any single sample.

Figure S3. Indicator species analysis examining associations between chemicals and land use types. Color indicates concentration and size the scaled frequency of occurrence. Significant associations are labeled with a black bar and the land use type they are associated with. No correction was made for multiple comparisons.

## References

Abbes, K., Biondi, A., Kurtulus, A., Ricupero, M., Russo, A., Siscaro, G., et al. (2015). Combined non-Target effects of insecticide and high temperature on the parasitoid bracon nigricans. PLoS One. doi: 10.1371/journal.pone.0138411.

Abivardi, C., Weber, D. C., and Dorn, S. (1999). Effects of carbaryl and cyhexatin on survival and reproductive behaviour of Cydia pomonella (Lepidoptera: Tortricidae). Ann. Appl. Biol. doi: 10.1111/j.1744-7348.1999.tb05250.x.

Ahmed, M. A. I., Temerak, S. A. H., Abdel-Galil, F. A. K., and Manna, S. H. M. (2016). Susceptibility of field and laboratory strains of Cotton leafworm, Spodoptera littoralis (Boisd.) (Lepidoptera: Noctuidae) to spinosad pesticide under laboratory conditions. Plant Prot. Sci. doi: 10.17221/5/2015-PPS.

Atwood, D., and Paisley-Jones, C. (2017). Pesticides Industry Sales and Usage 2008-2012. Mark. Estim.

Balbuena, M. S., Tison, L., Hahn, M. L., Greggers, U., Menzel, R., and Farina, W. M. (2015). Effects of sublethal doses of glyphosate on honeybee navigation. J. Exp. Biol. doi: 10.1242/jeb.117291.

Belsky, J., and Joshi, N. K. (2018). Assessing role of major drivers in recent decline of monarch butterfly population in North America. Front. Environ. Sci. doi: 10.3389/fenvs.2018.00086.

Bonmatin, J. M., Giorio, C., Girolami, V., Goulson, D., Kreutzweiser, D. P., Krupke, C., et al. (2015). Environmental fate and exposure; neonicotinoids and fipronil. Environ. Sci. Pollut. Res. 22, 35–67. doi: 10.1007/s11356-014-3332-7.

Brook, B. W., Sodhi, N. S., and Bradshaw, C. J. A. (2008). Synergies among extinction drivers under global change. Trends Ecol. Evol. doi: 10.1016/j.tree.2008.03.011.

Chen, J., Jiang, W., Hu, H., Ma, X., Li, Q., Song, X., et al. (2019). Joint toxicity of methoxyfenozide and lufenuron on larvae of Spodoptera exigua Hübner (Lepidoptera: Noctuidae). J. Asia. Pac. Entomol. doi: 10.1016/j.aspen.2019.06.004.

Crone, E. E., Pelton, E. M., Brown, L. M., Thomas, C. C., and Schultz, C. B. (2019). Why are monarch butterflies declining in the West? Understanding the importance of multiple correlated drivers. Ecol. Appl. doi: 10.1002/eap.1975.

Dai, P., Yan, Z., Ma, S., Yang, Y., Wang, Q., Hou, C., et al. (2018). The Herbicide Glyphosate Negatively Affects Midgut Bacterial Communities and Survival of Honey Bee during Larvae Reared in Vitro. J. Agric. Food Chem. doi: 10.1021/acs.jafc.8b02212.

David G. James (2019). A Neonicotinoid Insecticide at a Rate Found in Nectar Reduces Longevity but Not Oogenesis in Monarch. Insects.

De Cáceres, M., Jansen, F., and Dell, N. (2020). indicspecies: Relationship Between Species and Groups of Sites. Available at: https://cran.r-project.org/package=indicspecies.

De Cáceres, M., and Legendre, P. (2009). Associations between species and groups of sites: Indices and statistical inference. Ecology. doi: 10.1890/08-1823.1.

Deutsch, C. A., Tewksbury, J. J., Huey, R. B., Sheldon, K. S., Ghalambor, C. K., Haak, D. C., et al. (2008). Impacts of climate warming on terrestrial ectotherms across latitude. Proc. Natl. Acad. Sci. U. S. A. 105, 6668–72. doi: 10.1073/pnas.0709472105.

Egan, J. F., Bohnenblust, E., Goslee, S., Mortensen, D., and Tooker, J. (2014). Herbicide drift can affect plant and arthropod communities. Agric. Ecosyst. Environ. doi: 10.1016/j.agee.2013.12.017.

Epstein, L. (2014). Fifty Years Since Silent Spring. Annu. Rev. Phytopathol. doi: 10.1146/annurev-phyto-102313-045900.

Espeset, A. E., Harrison, J. G., Shapiro, A. M., Nice, C. C., Thorne, J. H., Waetjen, D. P., et al. (2016). Understanding a migratory species in a changing world: climatic effects and demographic declines in the western monarch revealed by four decades of intensive monitoring. Oecologia. doi: 10.1007/s00442-016-3600-y.

European Committee for Stantarization (2008). Foods of plant origin-Determination of pesticide residues using GC-MS and/or LC-MS/MS following acetonitrile extraction/partitioning and cleanup by dispersive SPE—QuEChERS-method. doi: 10.1038/s41598-017-05299-9.

Fantke, P., Gillespie, B. W., Juraske, R., and Jolliet, O. (2014). Estimating half-lives for pesticide dissipation from plants. Environ. Sci. Technol. doi: 10.1021/es500434p.

Fantke, P., and Juraske, R. (2013). Variability of pesticide dissipation half-lives in plants. Environ. Sci. Technol. doi: 10.1021/es303525x.

Forister, M. L., Cousens, B., Harrison, J. G., Anderson, K., Thorne, J. H., Waetjen, D., et al. (2016). Increasing neonicotinoid use and the declining butterfly fauna of lowland California. Biol. Lett. 12. doi: 10.1098/rsbl.2016.0475.

Forister, M. L., McCall, A. C., Sanders, N. J., Fordyce, J. A., Thorne, J. H., O’Brien, J., et al. (2010). Compounded effects of climate change and habitat alteration shift patterns of butterfly diversity. Proc. Natl. Acad. Sci. 107, 2088–2092. doi: 10.1073/pnas.0909686107.

Forister, M. L., Pelton, E. M., and Black, S. H. (2019). Declines in insect abundance and diversity: We know enough to act now. Conserv. Sci. Pract. doi: 10.1111/csp2.80.

Gibbs, K. E., MacKey, R. L., and Currie, D. J. (2009). Human land use, agriculture, pesticides and losses of imperiled species. Divers. Distrib. doi: 10.1111/j.1472-4642.2008.00543.x.

Gilburn, A. S., Bunnefeld, N., Wilson, J. M., Botham, M. S., Brereton, T. M., Fox, R., et al. (2015). Are neonicotinoid insecticides driving declines of widespread butterflies? PeerJ 3, e1402. doi: 10.7717/peerj.1402.

González-Varo, J. P., Biesmeijer, J. C., Bommarco, R., Potts, S. G., Schweiger, O., Smith, H. G., et al. (2013). Combined effects of global change pressures on animal-mediated pollination. Trends Ecol. Evol. doi: 10.1016/j.tree.2013.05.008.

Goulson, D., Nicholls, E., Botías, C., and Rotheray, E. L. (2015). Bee declines driven by combined Stress from parasites, pesticides, and lack of flowers. Science 347. doi: 10.1126/science.1255957.

Hallmann, C. A., Sorg, M., Jongejans, E., Siepel, H., Hofland, N., Schwan, H., et al. (2017). More than 75 percent decline over 27 years in total flying insect biomass in protected areas. PLoS One. doi: 10.1371/journal.pone.0185809.

Hill, M. O. (1973). Diversity and Evenness: A Unifying Notation and Its Consequences. Ecology. doi: 10.2307/1934352.

Hummelbrunner, L. A., and Isman, M. B. (2001). Acute, sublethal, antifeedant, and synergistic effects of monoterpenoid essential oil compounds on the tobacco cutworm, Spodoptera litura (Lep., Noctuidae). J. Agric. Food Chem. doi: 10.1021/jf000749t.

Janzen, D. H., and Hallwachs, W. (2019). Perspective: Where might be many tropical insects? Biol. Conserv. doi: 10.1016/j.biocon.2019.02.030.

Jones, M. M., Robertson, J. L., and Weinzierl, R. A. (2012a). Toxicity of Thiamethoxam and Mixtures of Chlorantraniliprole Plus Acetamiprid, Esfenvalerate, or Thiamethoxam to Neonates of Oriental Fruit Moth (Lepidoptera: Tortricidae). J. Econ. Entomol. doi: 10.1603/ec11349.

Jones, M. M., Robertson, J. L., and Weinzierl, R. A. (2012b). Toxicity of Thiamethoxam and Mixtures of Chlorantraniliprole Plus Acetamiprid, Esfenvalerate, or Thiamethoxam to Neonates of Oriental Fruit Moth (Lepidoptera: Tortricidae). J. Econ. Entomol. 105, 1426–1431. doi: 10.1603/ec11349.

Jost, L. (2006). Entropy and diversity. Oikos. doi: 10.1111/j.2006.0030-1299.14714.x.

Krischik, V., Rogers, M., Gupta, G., and Varshney, A. (2015). Soil-applied imidacloprid translocates to ornamental flowers and reduces survival of adult coleomegilla maculata, harmonia axyridis, and hippodamia convergens lady beetles, and larval danaus plexippus and vanessa cardui butterflies. PLoS One 10. doi: 10.1371/journal.pone.0119133.

Krishnan, N., Zhang, Y., Bidne, K. G., Hellmich, R. L., Coats, J. R., and Bradbury, S. P. (2020). Assessing field-scale risks of foliar insecticide applications to monarch butterfly (Danaus plexippus) larvae. Environ. Toxicol. Chem. doi: 10.1002/etc.4672.

Krupke, C. H., Hunt, G. J., Eitzer, B. D., Andino, G., and Given, K. (2012). Multiple routes of pesticide exposure for honey bees living near agricultural fields. PLoS One. doi: 10.1371/journal.pone.0029268.

LaLone, C., Villeneuve, D., Kristina, G., Tollefsen, K., and Ankley, G. (2014). Conservation of lepidopteran ecdysteroid receptor provides evidence for butterfly susceptibility to diacylhydrazine and bisacylhydrazine chemicals. in Society of Environmental Toxicology and Chemistry (Vancouver).

Legendre, P., and Legendre, L. (2012). Numerical ecology. Developments in environmental modeling.

Liu, Y., Zhang, H., He, F., Li, X., Tan, H., and Zeng, D. (2018a). Combined toxicity of chlorantraniliprole, lambda-cyhalothrin, and imidacloprid to the silkworm Bombyx mori (Lepidoptera: Bombycidae). Environ. Sci. Pollut. Res. doi: 10.1007/s11356-018-2374-7.

Liu, Y., Zhang, H., He, F., Li, X., Tan, H., and Zeng, D. (2018b). Combined toxicity of chlorantraniliprole, lambda-cyhalothrin, and imidacloprid to the silkworm Bombyx mori (Lepidoptera: Bombycidae). Environ. Sci. Pollut. Res. 25, 22598–22605. doi: 10.1007/s11356-018-2374-7.

Morrissey, C. A., Mineau, P., Devries, J. H., Sanchez-Bayo, F., Liess, M., Cavallaro, M. C., et al. (2015). Neonicotinoid contamination of global surface waters and associated risk to aquatic invertebrates: A review. Environ. Int. doi: 10.1016/j.envint.2014.10.024.

Motta, E. V. S., Raymann, K., and Moran, N. A. (2018). Glyphosate perturbs the gut microbiota of honey bees. Proc. Natl. Acad. Sci. U. S. A. doi: 10.1073/pnas.1803880115.

Mulé, R., Sabella, G., Robba, L., and Manachini, B. (2017). Systematic review of the effects of chemical insecticides on four common butterfly families. Front. Environ. Sci. doi: 10.3389/fenvs.2017.00032.

Nasr, H. M., Badawy, M. E. I., and Rabea, E. I. (2010). Toxicity and biochemical study of two insect growth regulators, buprofezin and pyriproxyfen, on cotton leafworm Spodoptera littoralis. Pestic. Biochem. Physiol. doi: 10.1016/j.pestbp.2010.06.007.

Nice, C. C., Forister, M. L., Harrison, J. G., Gompert, Z., Fordyce, J. A., Thorne, J. H., et al. (2019). Extreme heterogeneity of population response to climatic variation and the limits of prediction. Glob. Chang. Biol. doi: 10.1111/gcb.14593.

Oksanen, J., Blanchet, F. G., Friendly, M., Kindt, R., Legendre, P., McGlinn, D., et al. (2019). vegan: Community Ecology Package. Available at: https://cran.r-project.org/package=vegan.

Olaya-Arenas, P., and Kaplan, I. (2019). Quantifying pesticide exposure risk for monarch caterpillars on milkweeds bordering agricultural land. Front. Ecol. Evol. doi: 10.3389/fevo.2019.00223.

Pecenka, J. R., and Lundgren, J. G. (2015). Non-target effects of clothianidin on monarch butterflies. Sci. Nat. 102. doi: 10.1007/s00114-015-1270-y.

Pelton, E. M., Schultz, C. B., Jepsen, S. J., Black, S. H., and Crone, E. E. (2019). Western Monarch Population Plummets: Status, Probable Causes, and Recommended Conservation Actions. Front. Ecol. Evol. doi: 10.3389/fevo.2019.00258.

Perveen, F. (2000). Sublethal effects of chlorfluazuron on reproductivity and viability of Spodoptera litura (F.) (Lep., Noctuidae). J. Appl. Entomol. doi: 10.1046/j.1439-0418.2000.00468.x.

Pisa, L. W., Amaral-Rogers, V., Belzunces, L. P., Bonmatin, J. M., Downs, C. A., Goulson, D., et al. (2014). Effects of neonicotinoids and fipronil on non-target invertebrates. Environ. Sci. Pollut. Res. doi: 10.1007/s11356-014-3471-x.

Pocius, V. M., Debinski, D. M., Pleasants, J. M., Bidne, K. G., and Hellmich, R. L. (2018). Monarch butterflies do not place all of their eggs in one basket: Oviposition on nine Midwestern milkweed species. Ecosphere. doi: 10.1002/ecs2.2064.

Potts, S. G., Biesmeijer, J. C., Kremen, C., Neumann, P., Schweiger, O., and Kunin, W. E. (2010). Global pollinator declines: Trends, impacts and drivers. Trends Ecol. Evol. doi: 10.1016/j.tree.2010.01.007.

Rehan, A., and Freed, S. (2015). Fitness Cost of Methoxyfenozide and the Effects of Its Sublethal Doses on Development, Reproduction, and Survival of Spodoptera litura (Fabricius) (Lepidoptera: Noctuidae). Neotrop. Entomol. doi: 10.1007/s13744-015-0306-5.

Reiter, M. E., Wolder, M. A., Isola, J. E., Jongsomjit, D., Hickey, C. M., Carpenter, M., et al. (2015). Local and landscape habitat associations of shorebirds in wetlands of the Sacramento Valley of California. J. Fish Wildl. Manag. doi: 10.3996/012014-JFWM-003.

Sanchez-Bayo, F., and Goka, K. (2014). Pesticide residues and bees - A risk assessment. PLoS One. doi: 10.1371/journal.pone.0094482.

Sánchez-Bayo, F., and Wyckhuys, K. A. G. (2019). Worldwide decline of the entomofauna: A review of its drivers. Biol. Conserv. doi: 10.1016/j.biocon.2019.01.020.

Schubert, A., and Telcs, A. (2014). A note on the Jaccardized Czekanowski similarity index. Scientometrics. doi: 10.1007/s11192-013-1044-2.

Schultz, C. B., Brown, L. M., Pelton, E., and Crone, E. E. (2017). Citizen science monitoring demonstrates dramatic declines of monarch butterflies in western North America. Biol. Conserv. doi: 10.1016/j.biocon.2017.08.019.

Stenoien, C., Nail, K. R., Zalucki, J. M., Parry, H., Oberhauser, K. S., and Zalucki, M. P. (2018). Monarchs in decline: a collateral landscape-level effect of modern agriculture. Insect Sci. doi: 10.1111/1744-7917.12404.

Stoner, K. A., Cowles, R. S., Nurse, A., and Eitzer, B. D. (2019). Tracking Pesticide Residues to a Plant Genus Using Palynology in Pollen Trapped from Honey Bees (Hymenoptera: Apidae) at Ornamental Plant Nurseries. Environ. Entomol. doi: 10.1093/ee/nvz007.

Theobald, D. M. (2005). Landscape patterns of exurban growth in the USA from 1980 to 2020. Ecol. Soc. doi: 10.5751/ES-01390-100132.

Tylianakis, J. M., Didham, R. K., Bascompte, J., and Wardle, D. A. (2008). Global change and species interactions in terrestrial ecosystems. Ecol. Lett. doi: 10.1111/j.1461-0248.2008.01250.x.

Wagner, S. (2019). Study 310: Surface Water Monitoring for Pesticides in Agricultural Areas of Northern California, 2019. Sacramento.

Wang, D., Gong, P., Li, M., Qiu, X., and Wang, K. (2009). Sublethal effects of spinosad on survival, growth and reproduction of Helicoverpa armigera (Lepidoptera: Noctuidae). Pest Manag. Sci. doi: 10.1002/ps.1672.

Wang, D., Wang, Y.-M., Liu, H.-Y., Xin, Z., and Xue, M. (2013). Lethal and Sublethal Effects of Spinosad on <I>Spodoptera exigua</I> (Lepidoptera: Noctuidae). J. Econ. Entomol. doi: 10.1603/ec12220.

Wepprich, T., Adrion, J. R., Ries, L., Wiedmann, J., and Haddad, N. M. (2019). Butterfly abundance declines over 20 years of systematic monitoring in Ohio, USA. PLoS One. doi: 10.1371/journal.pone.0216270.

Weston, D. P., Holmes, R. W., and Lydy, M. J. (2009). Residential runoff as a source of pyrethroid pesticides to urban creeks. Environ. Pollut. doi: 10.1016/j.envpol.2008.06.037.

Whitehorn, P. R., O’connor, S., Wackers, F. L., and Goulson, D. (2012). Neonicotinoid pesticide reduces bumble bee colony growth and queen production lab field week cumulative weight gain (g). Science. doi: 10.5061/dryad.1805c973.

Wolfenbarger, L. L. R., Naranjo, S. E., Lundgren, J. G., Bitzer, R. J., and Watrud, L. S. (2008). Bt crop effects on functional guilds of non-target arthropods: A meta-analysis. PLoS One. doi: 10.1371/journal.pone.0002118.

Wood, T. J., and Goulson, D. (2017). The environmental risks of neonicotinoid pesticides: a review of the evidence post 2013. Environ. Sci. Pollut. Res. doi: 10.1007/s11356-017-9240-x.

Yue, B., Wilde, G. E., and Arthur, F. (2003). Evaluation of Thiamethoxam and Imidacloprid as Seed Treatments to Control European Corn Borer and Indianmeal Moth (Lepidoptera: Pyralidae) Larvae. J. Econ. Entomol. 96, 503–509. doi: 10.1093/jee/96.2.503.

Zhu, W., Schmehl, D. R., Mullin, C. A., and Frazier, J. L. (2014). Four common pesticides, their mixtures and a formulation solvent in the hive environment have high oral toxicity to honey bee larvae. PLoS One. doi: 10.1371/journal.pone.0077547.

