## Supplemental Information for "Pesticide contamination of milkweeds across the agricultural, urban, and open spaces of low elevation Northern California"

### Contents:

Table S1. Number of samples of different milkweed species from different land use types.

Table S2. Retention times and optimized SRM acquisition parameters for pesticides and internal standards (RT: Retention time, CE: Collision Energy)

Table S3. Contact and oral LD_50_ data for honeybees.

Table S4. Studies from Lepidoptera literature review.

Table S5. Exceedances of honeybee LD_50_ concentrations by land use type and compound.

Table S6. Indicator species results for associations between sites and individual compounds. Values in each land type category show mean concentration (ppb).

Figure S1. Mean effective numbers of pesticides per sample by land use type using different hill numbers after rarefaction. Points show the mean effective number of compounds per sample. Error bars show the range of effective numbers of pesticides across samples within one land use type.

### Figure S2. Variation in the number of compounds per sample by milkweed species. Bars show the maximum and minimum number of compounds detected in any single sample.

Table S1.Number of samples of different milkweed species from different land use types.

| *Asclepias* sp. | Agriculture | Refuge | Retail | Urban |
| --- | --- | --- | --- | --- |
| *Asclepias curassavica* | 0 | 0 | 9 | 3 |
| *Asclepias eriocarpa* | 4 | 0 | 0 | 0 |
| *Asclepias fasicularis* | 64 | 86 | 0 | 11 |
| *Asclepias speciosa* | 32 | 3 | 2 | 13 |

Table S2. Retention times and optimized SRM acquisition parameters for pesticides and internal standards (RT: Retention time, CE: Collision Energy)

| **Compound** | **RT (min)** | **Polarity** | **Precursor (m/z)** | **RF Lens (V)** | **Product 1 (m/z)** | **CE 1 (V)** | **Product 2 (m/z)** | **CE 2 (V)** |
| --- | --- | --- | --- | --- | --- | --- | --- | --- |
| d7-Propamocarb | 3.25 | Positive | 196.2 | 97 | 103.1 | 18 | 151.2 | 14 |
| d4-Imidacloprid | 5.05 | Positive | 260.1 | 114 | 213.1 | 16 | 179.1 | 19 |
| d7-Bentazone | 6.42 | Negative | 246.0 | 134 | 132.1 | 26 | 182.2 | 20 |
| d5-Atrazine | 7.42 | Positive | 221.0 | 113 | 179.1 | 18 | 101.0 | 25 |
| d10-Chlorpyrifos | 9.66 | Positive | 359.9 | 123 | 199.0 | 199 | 98.9 | 31 |
| Chlormequat chloride | 0.73 | Positive | 122.0 | 113 | 58.0 | 27 | 63.0 | 21 |
| Mepiquat chloride | 0.79 | Positive | 114.1 | 128 | 98.0 | 26 | 58.0 | 25 |
| Methamidophos | 1.79 | Positive | 141.9 | 100 | 94.0 | 14 | 125.0 | 14 |
| Cyromazine | 2.40 | Positive | 167.0 | 133 | 85.1 | 19 | 125.1 | 18 |
| Acephate | 2.74 | Positive | 184.0 | 65 | 143.0 | 10 | 94.8 | 23 |
| Omethoate | 3.24 | Positive | 214.0 | 115 | 182.8 | 10 | 124.9 | 18 |
| Propamocarb | 3.28 | Positive | 189.1 | 98 | 102.0 | 17 | 74.1 | 25 |
| Aminocarb | 3.32 | Positive | 209.1 | 124 | 137.1 | 24 | 152.1 | 14 |
| Formetanate hydrochloride | 3.35 | Positive | 222.1 | 144 | 165.1 | 15 | 120 | 27 |
| Butocarboxim sulfoxide | 3.47 | Positive | 207.0 | 94 | 132.0 | 10 | 88.0 | 10 |
| Pymetrozine | 3.57 | Positive | 218.0 | 150 | 105.0 | 20 | 78.1 | 39 |
| Dinotefuran | 3.59 | Positive | 203.0 | 98 | 113.1 | 10 | 129.1 | 12 |
| Butoxycarboxim | 3.65 | Positive | 223.0 | 124 | 166.1 | 15 | 46.1 | 26 |
| Aldicarb sulfone | 3.71 | Positive | 223.1 | 117 | 148.0 | 10 | 86.1 | 16 |
| Oxamyl | 3.90 | Positive | 237.0 | 73 | 72.1 | 10 | 90.0 | 10 |
| Methomyl | 4.13 | Positive | 163.1 | 71 | 87.9 | 10 | 106.1 | 10 |
| Demeton-S-methylsulfone | 4.25 | Positive | 263.0 | 150 | 169.0 | 16 | 108.9 | 28 |
| Thiamethoxam | 4.39 | Positive | 292.0 | 121 | 211.1 | 12 | 181.0 | 22 |
| Mexacarbate | 4.50 | Positive | 223.1 | 136 | 151.1 | 24 | 166.1 | 15 |
| Monocrotophos | 4.54 | Positive | 224.0 | 112 | 127.0 | 16 | 192.9 | 10 |
| Ethiofencarb sulfone | 4.73 | Positive | 258.0 | 119 | 107.0 | 16 | 201.0 | 10 |
| Dicrotophos | 4.81 | Positive | 238.0 | 127 | 127.0 | 18 | 192.5 | 10 |
| Pirimicarb-desmethyl | 4.84 | Positive | 225.2 | 135 | 168.0 | 15 | 72.0 | 21 |
| Ethiofencarb sulfoxide | 4.91 | Positive | 242.0 | 106 | 107.0 | 18 | 185.0 | 10 |
| Trichlorfon | 4.94 | Positive | 256.9 | 131 | 108.9 | 18 | 79.0 | 30 |
| Clothianidin | 5.03 | Positive | 250.0 | 104 | 169.0 | 13 | 131.9 | 17 |
| Imidacloprid | 5.06 | Positive | 256.0 | 131 | 209.0 | 16 | 175.1 | 19 |
| Fenuron | 5.13 | Positive | 164.8 | 116 | 72.1 | 15 | 46.1 | 15 |
| Thiabendazole | 5.13 | Positive | 202.0 | 208 | 175.0 | 26 | 131.0 | 33 |
| Flumetsulam | 5.17 | Positive | 326.0 | 192 | 129.0 | 26 | 262.1 | 19 |
| Dimethoate | 5.19 | Positive | 229.8 | 106 | 198.8 | 10 | 124.9 | 22 |
| 3-Hydroxy-carbofuran | 5.21 | Positive | 238.1 | 119 | 181.0 | 10 | 163.1 | 16 |
| Vamidothion | 5.25 | Positive | 288.0 | 120 | 146.0 | 14 | 118.1 | 23 |
| Fuberidazole | 5.26 | Positive | 185.1 | 152 | 157.1 | 22 | 129.0 | 35 |
| Metamitron | 5.29 | Positive | 203.0 | 170 | 174.8 | 17 | 104.0 | 23 |
| Methiocarb sulfoxide | 5.29 | Positive | 242.0 | 134 | 185.0 | 14 | 122.1 | 29 |
| Chloridazon | 5.41 | Positive | 222.0 | 152 | 104.0 | 23 | 92.0 | 26 |
| Acetamiprid | 5.55 | Positive | 223.0 | 118 | 126.0 | 21 | 90.0 | 34 |
| Methiocarb sulfone | 5.62 | Positive | 258.0 | 119 | 122.0 | 19 | 201.0 | 10 |
| Schradan | 5.69 | Positive | 287.1 | 144 | 242.1 | 14 | 135.1 | 26 |
| Mevinphos | 5.83 | Positive | 225.0 | 97 | 127.0 | 17 | 192.9 | 10 |
| Ethirimol | 5.89 | Positive | 209.8 | 192 | 140.1 | 22 | 98.0 | 27 |
| Florasulam | 5.96 | Positive | 360.0 | 179 | 129.0 | 25 | 108.9 | 53 |
| Pirimicarb | 5.97 | Positive | 239.1 | 147 | 182.1 | 16 | 72.0 | 21 |
| Thiacloprid | 5.98 | Positive | 253.0 | 162 | 126.0 | 21 | 90.0 | 36 |
| Metoxuron | 6.21 | Positive | 229.0 | 145 | 72.1 | 18 | 156.0 | 26 |
| Formothion | 6.23 | Positive | 258.0 | 80 | 199.0 | 10 | 124.9 | 22 |
| Imazethapyr | 6.29 | Positive | 290.1 | 189 | 177.0 | 27 | 248.1 | 19 |
| Carbetamide | 6.33 | Positive | 237.1 | 102 | 192.0 | 10 | 120.0 | 16 |
| Metolcarb | 6.34 | Positive | 166.0 | 83 | 108.9 | 10 | 94.0 | 31 |
| Oxadixyl | 6.35 | Positive | 279.1 | 120 | 219.1 | 10 | 132.1 | 31 |
| Tricyclazole | 6.45 | Positive | 190.0 | 178 | 163.0 | 23 | 136.0 | 29 |
| Bentazone | 6.46 | Negative | 238.9 | 169 | 132.0 | 26 | 197.0 | 21 |
| Cyanazine | 6.46 | Positive | 241.1 | 164 | 214.1 | 18 | 103.9 | 29 |
| Azamethiphos | 6.56 | Positive | 324.9 | 166 | 182.9 | 16 | 111.9 | 34 |
| Bromacil | 6.60 | Positive | 261.0 | 112 | 204.9 | 14 | 187.8 | 28 |
| Propoxur | 6.60 | Positive | 210.0 | 87 | 111.0 | 14 | 168.1 | 10 |
| Thiophanate-methyl | 6.62 | Positive | 343.0 | 161 | 151.0 | 20 | 93.0 | 46 |
| Bendiocarb | 6.67 | Positive | 224.0 | 104 | 167.1 | 10 | 108.9 | 18 |
| Carbofuran | 6.68 | Positive | 222.0 | 111 | 165.1 | 12 | 123.0 | 22 |
| Ofurace | 6.76 | Positive | 282.1 | 145 | 254.1 | 12 | 160.1 | 24 |
| Malaoxon | 6.79 | Positive | 315.0 | 133 | 98.9 | 23 | 269.0 | 10 |
| Imazaquin | 6.81 | Positive | 312.1 | 195 | 267.1 | 21 | 199.0 | 28 |
| Thidiazuron | 6.83 | Positive | 220.6 | 119 | 101.9 | 16 | 127.9 | 17 |
| Pyroxsulam | 6.83 | Positive | 435.0 | 262 | 195.1 | 26 | 258.0 | 22 |
| Simetryn | 6.83 | Positive | 214.1 | 161 | 124.1 | 20 | 96.0 | 25 |
| Desmetryn | 6.85 | Positive | 214.1 | 161 | 172.0 | 18 | 82.0 | 30 |
| Ancymidol | 6.86 | Positive | 257.1 | 157 | 135.0 | 25 | 81.1 | 25 |
| Hexazinone | 6.89 | Positive | 253.1 | 141 | 171.1 | 16 | 71.1 | 31 |
| Tebuthiuron | 6.90 | Positive | 229.0 | 145 | 172.1 | 18 | 116.0 | 27 |
| Metosulam | 6.99 | Positive | 418.0 | 233 | 174.9 | 27 | 140.0 | 50 |
| Prometon | 7.08 | Positive | 226.1 | 156 | 184.1 | 19 | 142.1 | 23 |
| Carbaryl | 7.08 | Positive | 202.0 | 95 | 145.1 | 10 | 127.0 | 29 |
| Fenthion sulfoxide | 7.09 | Positive | 295.0 | 187 | 280.0 | 19 | 108.9 | 32 |
| Ethiofencarb | 7.09 | Positive | 226.1 | 105 | 107.0 | 17 | 164.1 | 10 |
| Cyantraniliprole | 7.10 | Positive | 475.0 | 158 | 285.9 | 11 | 444.0 | 18 |
| Terbumeton | 7.20 | Positive | 226.2 | 153 | 170.0 | 17 | 142.1 | 23 |
| Monolinuron | 7.20 | Positive | 215.0 | 131 | 126.0 | 18 | 148.0 | 15 |
| Fosthiazate | 7.21 | Positive | 284.0 | 118 | 103.9 | 21 | 228.0 | 10 |
| Fluometuron | 7.24 | Positive | 233.0 | 145 | 72.0 | 19 | 46 | 18 |
| 2,4-D | 7.27 | Negative | 218.9 | 101 | 160.9 | 13 | 125.0 | 26 |
| Bromoxynil | 7.29 | Negative | 275.8 | 194 | 80.9 | 31 | 78.9 | 30 |
| DNOC | 7.30 | Negative | 197.0 | 147 | 180.0 | 19 | 137.0 | 18 |
| Ethoxyquin | 7.36 | Positive | 218.1 | 183 | 160.1 | 33 | 148.1 | 22 |
| Benodanil | 7.36 | Positive | 323.8 | 180 | 231.0 | 23 | 202.9 | 35 |
| Imazalil | 7.37 | Positive | 297.0 | 170 | 156.0 | 23 | 200.9 | 18 |
| Isoprocarb | 7.38 | Positive | 194.1 | 105 | 95.0 | 15 | 137.1 | 10 |
| Flutriafol | 7.42 | Positive | 302.0 | 144 | 70.0 | 19 | 123.0 | 28 |
| Chlorotoluron | 7.43 | Positive | 213.0 | 141 | 72.1 | 18 | 46.0 | 16 |
| Atrazine | 7.44 | Positive | 216.1 | 167 | 174.0 | 18 | 103.9 | 28 |
| Metobromuron | 7.47 | Positive | 258.9 | 114 | 148.0 | 15 | 169.9 | 19 |
| Metazachlor | 7.48 | Positive | 278.0 | 111 | 210.1 | 10 | 134.1 | 22 |
| Lenacil | 7.50 | Positive | 235.1 | 109 | 153.1 | 16 | 136.0 | 32 |
| Isocarbophos | 7.53 | Positive | 307.0 | 73 | 231.0 | 16 | 273.0 | 10 |
| Metalxyl | 7.54 | Positive | 280.1 | 98 | 220.0 | 14 | 192.2 | 18 |
| Griseofulvin | 7.54 | Positive | 353.2 | 188 | 285.0 | 18 | 165.1 | 20 |
| Methoprotryne | 7.59 | Positive | 272.1 | 175 | 240.2 | 19 | 198.0 | 23 |
| Isoproturon | 7.59 | Positive | 207.1 | 143 | 72.1 | 19 | 165.1 | 14 |
| Fensulfothion | 7.64 | Positive | 309.0 | 180 | 280.9 | 15 | 253.0 | 18 |
| Heptenophos | 7.69 | Positive | 251.0 | 123 | 127.0 | 17 | 124.9 | 13 |
| Desmedipham | 7.71 | Positive | 301.3 | 133 | 182.0 | 10 | 136.0 | 20 |
| Forchlorfenuron | 7.71 | Positive | 248.0 | 134 | 129.0 | 18 | 93.0 | 33 |
| Dodemorph | 7.73 | Positive | 282.2 | 188 | 116.1 | 21 | 98.0 | 27 |
| Cycluron | 7.73 | Positive | 199.1 | 137 | 72.1 | 22 | 69.1 | 21 |
| Chlorantraniliprole | 7.75 | Positive | 481.9 | 182 | 283.9 | 12 | 450.8 | 18 |
| Methabenzthiazuron | 7.75 | Positive | 222.0 | 118 | 165.1 | 17 | 150.0 | 33 |
| Diuron | 7.77 | Positive | 233.0 | 145 | 72.1 | 19 | 46 | 18 |
| Ioxynil | 7.79 | Negative | 369.7 | 204 | 126.8 | 35 | 214.9 | 32 |
| Azaconazole | 7.82 | Positive | 299.9 | 162 | 159.0 | 28 | 231.0 | 17 |
| Phenmedipham | 7.82 | Positive | 301.1 | 145 | 168.0 | 10 | 136.0 | 20 |
| Dimefuron | 7.83 | Positive | 339.0 | 220 | 167.0 | 22 | 72.1 | 26 |
| Benoxacor | 7.84 | Positive | 260.0 | 173 | 149.1 | 18 | 134.0 | 29 |
| Clomazone | 7.91 | Positive | 240.0 | 134 | 125.0 | 21 | 89.0 | 47 |
| Diethofencarb | 7.92 | Positive | 268.0 | 114 | 226.1 | 10 | 124.0 | 32 |
| Azinphos-methyl | 7.93 | Positive | 317.9 | 103 | 132.0 | 15 | 125.0 | 17 |
| Fenobucarb | 7.93 | Positive | 208.1 | 110 | 95.0 | 15 | 152.0 | 10 |
| Ethofumesate | 7.98 | Positive | 287.1 | 159 | 121.0 | 16 | 259.1 | 10 |
| Fluazifop | 8.01 | Positive | 328.0 | 174 | 282.0 | 19 | 254.0 | 26 |
| Azoxystrobin | 8.03 | Positive | 404.1 | 175 | 372.0 | 14 | 344.1 | 25 |
| Propazine | 8.03 | Positive | 230.1 | 177 | 146.1 | 23 | 188.1 | 18 |
| Pyrimethanil | 8.03 | Positive | 200.1 | 184 | 107.0 | 25 | 168.1 | 30 |
| Nuarimol | 8.04 | Positive | 315.1 | 177 | 252.1 | 22 | 243.0 | 25 |
| Ethiprole | 8.04 | Positive | 396.9 | 189 | 350.9 | 21 | 255.0 | 36 |
| Fenamidone | 8.05 | Positive | 312.1 | 151 | 236.1 | 15 | 92.0 | 25 |
| Halofenozide | 8.07 | Positive | 331.0 | 99 | 275.1 | 10 | 105.0 | 18 |
| Dimethenamid | 8.09 | Positive | 276.1 | 135 | 244.1 | 14 | 168.1 | 24 |
| Prometryn | 8.11 | Positive | 242.2 | 149 | 158.0 | 24 | 200.0 | 19 |
| Methiocarb | 8.13 | Positive | 226.1 | 105 | 169.1 | 10 | 121.0 | 19 |
| Spiroxamine | 8.18 | Positive | 298.2 | 167 | 144.2 | 20 | 100.0 | 30 |
| Crotoxyphos | 8.18 | Positive | 332.0 | 100 | 210.9 | 10 | 127.0 | 25 |
| Mandipropamid | 8.18 | Positive | 412.1 | 186 | 328.1 | 15 | 356.1 | 10 |
| Terbuthylazine | 8.19 | Positive | 230.1 | 140 | 174.0 | 17 | 132.0 | 25 |
| Boscalid | 8.20 | Positive | 343.0 | 174 | 307.0 | 21 | 272.0 | 30 |
| Isoxaben | 8.20 | Positive | 333.2 | 167 | 164.9 | 19 | 150.0 | 39 |
| Promecarb | 8.21 | Positive | 208.1 | 107 | 109.0 | 16 | 151.1 | 10 |
| Paclobutrazol | 8.22 | Positive | 294.0 | 151 | 70.1 | 21 | 125.0 | 38 |
| Terbutryn | 8.23 | Positive | 242.1 | 158 | 186.0 | 19 | 91.0 | 28 |
| Fluopicolide | 8.23 | Positive | 382.9 | 193 | 172.9 | 23 | 144.9 | 48 |
| Propyzamide | 8.26 | Positive | 256.0 | 110 | 190.0 | 14 | 173.0 | 29 |
| Mepronil | 8.27 | Positive | 270.1 | 159 | 118.9 | 24 | 228.1 | 15 |
| Fluxapyroxad | 8.28 | Positive | 382.0 | 140 | 362.1 | 13 | 342.1 | 20 |
| Fludioxonil | 8.28 | Negative | 247.0 | 149 | 180.0 | 28 | 126.1 | 31 |
| Isoprothiolane | 8.32 | Positive | 291.1 | 116 | 231.0 | 10 | 188.8 | 22 |
| Methoxyfenozide | 8.32 | Positive | 369.2 | 113 | 149.1 | 17 | 313.1 | 10 |
| Dimethomorph | 8.33 | Positive | 388.1 | 225 | 301.0 | 21 | 165.1 | 32 |
| Triadimefon | 8.34 | Positive | 294.0 | 138 | 197.0 | 16 | 141.0 | 22 |
| Propetamphos | 8.34 | Positive | 282.0 | 106 | 138.0 | 17 | 156.0 | 10 |
| Myclobutanil | 8.39 | Positive | 289.0 | 121 | 70.1 | 18 | 124.9 | 33 |
| Fluorochloridone | 8.40 | Positive | 312.0 | 132 | 292.0 | 21 | 145.0 | 48 |
| Butafenacil | 8.42 | Positive | 492.1 | 170 | 331.0 | 24 | 349.0 | 15 |
| Cumyluron | 8.43 | Positive | 303.1 | 137 | 184.9 | 13 | 125.0 | 33 |
| Fluopyram | 8.43 | Positive | 397.0 | 202 | 208.0 | 22 | 173.0 | 29 |
| Iprovalicarb | 8.44 | Positive | 321.2 | 129 | 119.1 | 19 | 116.1 | 20 |
| Fenhexamid | 8.45 | Positive | 302.0 | 166 | 97.0 | 23 | 55.0 | 35 |
| Bifenazate | 8.46 | Positive | 301.1 | 119 | 198.0 | 10 | 170.1 | 19 |
| Fluoxastrobin | 8.49 | Positive | 459.1 | 219 | 427.0 | 17 | 188.0 | 35 |
| Triazophos | 8.49 | Positive | 314.0 | 164 | 162.1 | 19 | 119.0 | 34 |
| Mefenacet | 8.49 | Positive | 299.0 | 132 | 148.1 | 14 | 120.0 | 25 |
| Spirotetramat | 8.50 | Positive | 374.1 | 185 | 302.1 | 17 | 330.2 | 15 |
| Bupirimate | 8.50 | Positive | 317.1 | 204 | 166.1 | 21 | 272.1 | 20 |
| Fluquinoconazole | 8.50 | Positive | 375.9 | 130 | 349.0 | 19 | 307.1 | 26 |
| Flufenacet | 8.51 | Positive | 364.0 | 126 | 194.0 | 10 | 152.1 | 19 |
| Tepraloxydim | 8.52 | Positive | 342.1 | 153 | 250.2 | 13 | 166.1 | 21 |
| Simeconazole | 8.53 | Positive | 294.1 | 150 | 70.1 | 20 | 135.1 | 21 |
| Chloroxuron | 8.54 | Positive | 291.0 | 178 | 72.1 | 21 | 46.0 | 19 |
| Tetraconazole | 8.56 | Positive | 372.0 | 187 | 158.9 | 30 | 123.0 | 55 |
| Dimethametryn | 8.58 | Positive | 256.1 | 180 | 185.9 | 21 | 96.0 | 30 |
| Trietazine | 8.58 | Positive | 230.1 | 178 | 132.0 | 22 | 104.0 | 29 |
| Cyazofamid | 8.63 | Positive | 325.0 | 122 | 107.9 | 14 | 217.0 | 18 |
| Napropamide | 8.65 | Positive | 272.0 | 149 | 171.1 | 19 | 199.0 | 13 |
| Alachlor | 8.64 | Positive | 270.1 | 112 | 238.1 | 10 | 162.2 | 20 |
| Metolachlor | 8.64 | Positive | 284.1 | 143 | 252.1 | 15 | 176.1 | 26 |
| Fipronil | 8.65 | Negative | 434.9 | 138 | 330.0 | 15 | 250.0 | 26 |
| Epoxiconazole | 8.68 | Positive | 330.0 | 149 | 121.0 | 21 | 100.9 | 44 |
| Fenbuconazole | 8.70 | Positive | 337.1 | 188 | 125.0 | 31 | 70.1 | 21 |
| Fenamiphos | 8.72 | Positive | 304.1 | 180 | 217.0 | 18 | 201.9 | 35 |
| Haloxyfop | 8.72 | Positive | 361.8 | 166 | 316.0 | 18 | 91.0 | 30 |
| Picoxystrobin | 8.72 | Positive | 368.1 | 111 | 145.1 | 21 | 205.0 | 10 |
| Tebufenozide | 8.73 | Positive | 353.2 | 110 | 297.1 | 10 | 133.1 | 19 |
| Triadimenol | 8.73 | Positive | 297.0 | 101 | 133.0 | 14 | 105.0 | 39 |
| Flubendiamide | 8.74 | Positive | 683.0 | 195 | 407.9 | 10 | 274.0 | 30 |
| Rotenone | 8.74 | Positive | 395.1 | 231 | 213.1 | 23 | 192.1 | 24 |
| Fenoxycarb | 8.75 | Positive | 302.0 | 152 | 88.0 | 19 | 116.0 | 10 |
| Flusilazole | 8.76 | Positive | 316.1 | 192 | 247.1 | 18 | 165.1 | 27 |
| Carfentrazone-ethyl | 8.76 | Positive | 412.0 | 217 | 346.0 | 23 | 365.9 | 18 |
| Diflubenzuron | 8.76 | Positive | 311.0 | 131 | 158.0 | 13 | 141.1 | 32 |
| Dimoxystrobin | 8.77 | Positive | 327.2 | 108 | 204.9 | 10 | 115.9 | 22 |
| Phenthoate | 8.78 | Positive | 320.8 | 119 | 247.1 | 12 | 135.0 | 20 |
| Isoxadifen-ethyl | 8.79 | Positive | 296.1 | 146 | 232.1 | 17 | 263.2 | 10 |
| Kresoxim-methyl | 8.79 | Positive | 314.1 | 116 | 267.1 | 10 | 222.1 | 13 |
| Neburon | 8.80 | Positive | 275.0 | 171 | 88.1 | 16 | 57.0 | 21 |
| Sulfotep | 8.81 | Positive | 323.0 | 145 | 171.0 | 14 | 114.9 | 29 |
| Penthiopyrad | 8.84 | Positive | 360.1 | 149 | 276.0 | 14 | 256.1 | 20 |
| Fipronil sulfone | 8.86 | Negative | 450.8 | 152 | 415.0 | 15 | 282.0 | 26 |
| Tebuconazole | 8.87 | Positive | 308.0 | 162 | 70.1 | 23 | 125.0 | 38 |
| Cyprodynil | 8.88 | Positive | 226.1 | 153 | 93.0 | 34 | 108.1 | 26 |
| Anilofos | 8.89 | Positive | 368.0 | 173 | 198.9 | 14 | 124.9 | 31 |
| Bromuconazole | 8.90 | Positive | 377.8 | 185 | 159.0 | 30 | 161.0 | 31 |
| Carpropamid | 8.90 | Positive | 334.0 | 126 | 139.0 | 20 | 195.9 | 13 |
| Etrimfos | 8.90 | Positive | 293.0 | 178 | 265.1 | 17 | 124.9 | 26 |
| Chlorfenvinphos | 8.93 | Positive | 359.0 | 159 | 155.1 | 13 | 169.9 | 39 |
| Penconazole | 8.93 | Positive | 284.1 | 134 | 159.0 | 30 | 70.1 | 18 |
| Zoxamide | 8.95 | Positive | 336.0 | 167 | 186.9 | 22 | 159.0 | 39 |
| Benzoylprop-ethyl | 8.95 | Positive | 366.0 | 134 | 105.0 | 16 | 77.1 | 48 |
| Fenthion | 8.96 | Positive | 279.0 | 133 | 169.0 | 16 | 247.0 | 13 |
| Cyflufenamid | 8.99 | Positive | 413.0 | 161 | 295.0 | 15 | 241.1 | 23 |
| Propiconazole | 8.99 | Positive | 342.1 | 69 | 159.0 | 30 | 122.9 | 55 |
| Pirimiphos-methyl | 9.00 | Positive | 306.1 | 199 | 164.1 | 22 | 108.0 | 31 |
| Coumaphos | 9.01 | Positive | 362.9 | 169 | 227.0 | 26 | 306.8 | 18 |
| Hexaconazole | 9.01 | Positive | 314.0 | 151 | 70.0 | 21 | 159.0 | 32 |
| Metconazole | 9.03 | Positive | 320.1 | 170 | 70.1 | 24 | 125.0 | 38 |
| Phoxim | 9.04 | Positive | 299.0 | 95 | 129.0 | 11 | 77.0 | 30 |
| Pyraclostrobin | 9.04 | Positive | 387.8 | 159 | 194.0 | 12 | 163.1 | 24 |
| Benzoximate | 9.06 | Positive | 364.1 | 105 | 199.0 | 10 | 105.0 | 24 |
| Prochloraz | 9.07 | Positive | 376.0 | 125 | 307.9 | 10 | 266.0 | 17 |
| Spinosad (Spinosyn A) | 9.09 | Positive | 732.4 | 299 | 142.1 | 29 | 98.1 | 45 |
| Metrafenone | 9.10 | Positive | 409.0 | 169 | 209.1 | 14 | 227.0 | 21 |
| Pencycuron | 9.12 | Positive | 329.1 | 190 | 125.0 | 40 | 218.1 | 16 |
| Haloxyfop-methyl | 9.15 | Positive | 376.0 | 180 | 315.8 | 17 | 91.0 | 31 |
| Thiobencarb | 9.15 | Positive | 258.0 | 118 | 125.0 | 20 | 89.0 | 49 |
| Indoxacarb | 9.16 | Positive | 528.0 | 233 | 203.0 | 38 | 150.0 | 24 |
| Diniconazole | 9.16 | Positive | 326.0 | 178 | 70.0 | 26 | 159.1 | 31 |
| Trifloxystrobin | 9.18 | Positive | 409.0 | 179 | 186.0 | 18 | 145.0 | 44 |
| Piperophos | 9.19 | Positive | 354.1 | 175 | 171.0 | 22 | 255.0 | 14 |
| Difenoconazole | 9.24 | Positive | 406.0 | 214 | 251.0 | 26 | 337.0 | 18 |
| Dithiopyr | 9.24 | Positive | 402.0 | 167 | 354.0 | 18 | 272.0 | 29 |
| Cycloate | 9.25 | Positive | 216.0 | 126 | 154.1 | 12 | 83.1 | 16 |
| Hexaflumuron | 9.28 | Negative | 458.8 | 142 | 438.9 | 10 | 175.0 | 36 |
| Clethodim | 9.27 | Positive | 360.1 | 140 | 164.1 | 18 | 268.1 | 12 |
| Prosulfocarb | 9.34 | Positive | 252.1 | 139 | 91.0 | 22 | 128.1 | 13 |
| Triflumizole | 9.34 | Positive | 346.0 | 113 | 278.1 | 10 | 73.1 | 17 |
| Furathiocarb | 9.38 | Positive | 383.1 | 176 | 194.9 | 18 | 252.1 | 13 |
| Quizalofop-ethyl | 9.39 | Positive | 373.0 | 221 | 299.0 | 19 | 271.1 | 26 |
| Buprofezin | 9.40 | Positive | 306.1 | 129 | 201.1 | 12 | 116.0 | 16 |
| Profenophos | 9.41 | Positive | 374.9 | 171 | 304.8 | 19 | 346.9 | 13 |
| Tetramethrin | 9.41 | Positive | 332.2 | 139 | 164.1 | 24 | 135.1 | 18 |
| Sethoxydim | 9.43 | Positive | 328.2 | 153 | 178.0 | 19 | 282.1 | 12 |
| Fluazinam | 9.44 | Negative | 462.8 | 194 | 415.9 | 19 | 398.0 | 16 |
| Tebufenpyrad | 9.44 | Positive | 334.2 | 206 | 117.0 | 36 | 145.0 | 27 |
| Esprocarb | 9.47 | Positive | 266.2 | 139 | 91.0 | 24 | 71.1 | 15 |
| Piperonyl butoxide | 9.51 | Positive | 356.3 | 118 | 177.1 | 10 | 119.0 | 33 |
| Tolfenpyrad | 9.58 | Positive | 384.1 | 188 | 197.0 | 25 | 196.0 | 19 |
| Imibenconazole | 9.54 | Positive | 411.0 | 209 | 125.0 | 31 | 171.0 | 10 |
| Hexythiazox | 9.63 | Positive | 353.1 | 123 | 228.0 | 15 | 168.0 | 25 |
| Tralkoxydim | 9.64 | Positive | 330.2 | 159 | 284.2 | 13 | 138.1 | 20 |
| Chlorpyrifos | 9.66 | Positive | 349.9 | 126 | 197.9 | 19 | 321.7 | 11 |
| Spiromesifen | 9.68 | Positive | 371.1 | 132 | 273.2 | 10 | 255.2 | 23 |
| Flufenoxuron | 9.69 | Positive | 489.2 | 142 | 158.1 | 18 | 140.9 | 41 |
| Sulprofos | 9.69 | Positive | 323.0 | 118 | 218.9 | 16 | 247.0 | 12 |
| Etoxazole | 9.74 | Positive | 360.1 | 201 | 141.0 | 31 | 304.0 | 18 |
| Quinoxyfen | 9.81 | Positive | 308.0 | 234 | 196.9 | 32 | 162.0 | 45 |
| Chlorfluazuron | 9.82 | Positive | 541.8 | 223 | 384.9 | 21 | 158.0 | 20 |
| Difenacoum | 9.84 | Positive | 445.1 | 236 | 179.0 | 31 | 257.2 | 20 |
| Amitraz | 9.87 | Positive | 294.1 | 103 | 163.1 | 14 | 122.1 | 29 |
| Fenpyroximate | 9.88 | Positive | 422.2 | 197 | 366.2 | 16 | 214.1 | 30 |
| Avermectin B1a | 9.95 | Positive | 890.5 | 225 | 305.1 | 25 | 567.2 | 14 |
| Resmethrin | 10.03 | Positive | 339.1 | 158 | 171.1 | 15 | 128.0 | 41 |
| Brodifacoum | 10.12 | Positive | 523.1 | 289 | 335.0 | 22 | 178.0 | 34 |
| Fenazaquin | 10.24 | Positive | 307.1 | 159 | 161.2 | 17 | 57.1 | 23 |
| Etofenprox | 10.25 | Positive | 394.0 | 133 | 177.1 | 15 | 359.2 | 10 |

Table S3. Contact and oral LD_50_ data for honey bees.

| Compound Name | Compound Type | Contact LD_50_ (ppb) | Oral LD_50_ (ppb) |
| --- | --- | --- | --- |
| Azoxystrobin | fungicide | 200000 | 25000 |
| Bifenazate | insecticide | 7800 | 98000 |
| Boscalid | fungicide | 200000 | 166000 |
| Buprofezin | insecticide | 200000 | 163500 |
| Carbaryl | insecticide | 378 | 210 |
| Carfentrazone ethyl | herbicide | 200000 | 200000 |
| Chlorantraniliprole | insecticide | 4000 | 104100 |
| Clethodim | herbicide | 51000 | 43000 |
| Clothianidin | insecticide | 44 | 4 |
| Cyantraniliprole | insecticide | 93 | 116 |
| Cyflufenamid | fungicide | 100000 | 100000 |
| Cyprodinil | fungicide | 784000 | 112500 |
| Cyromazine | insecticide | 200000 | 186000 |
| Difenoconazole | fungicide | 100000 | 177000 |
| Diflubenzuron | insecticide | 114800 | 25000 |
| Dinotefuran | insecticide | 41 | 18 |
| Diuron | herbicide | 101700 | 86750 |
| Etofenprox | insecticide | 38 | 366 |
| Etoxazole | insecticide | 200000 | 200000 |
| Fenhexamid | fungicide | 200000 | 102070 |
| Fenpyroximate | insecticide | 15800 | 118500 |
| Fipronil | insecticide | 6 | 4 |
| Fludioxonil | fungicide | 100000 | 100000 |
| Fluopicolide | fungicide | 100000 | 241000 |
| Fluopyram | fungicide | 100000 | 102300 |
| Fluoxastrobin | fungicide | 200000 | 843000 |
| Fluxapyroxad | fungicide | 100000 | 110900 |
| Hexythiazox | insecticide | 200000 | 112000 |
| Imazalil | fungicide | 39000 | 35100 |
| Imidacloprid | insecticide | 65 | 4 |
| Isoprothiolane | fungicide | NA | NA |
| Isoxaben | herbicide | 100000 | 100000 |
| Mepiquat | herbicide | 100000 | 107400 |
| Metalaxyl | fungicide | 200000 | 269000 |
| Methamidophos | insecticide | 1370 | 860 |
| Methiocarb | insecticide | 294 | 80 |
| Methoxyfenozide | insecticide | 100000 | 100000 |
| Metolachlor | herbicide | 110000 | 110000 |
| Metrafenone | fungicide | 100000 | 114000 |
| Myclobutanil | fungicide | 33900 | 33900 |
| Paclobutrazole | herbicide | 40000 | 2000 |
| Penthiopyrad | fungicide | 500000 | 500000 |
| Piperonyl butoxide | synergist | 294000 | NA |
| Prometryn | herbicide | 99000 | NA |
| Propamocarb | fungicide | 100000 | 84000 |
| Propiconazole | fungicide | 100000 | 100000 |
| Propyzamide | herbicide | 136000 | 100000 |
| Pyraclostrobin | fungicide | 100000 | 110000 |
| Spinosyn | insecticide | 47 | 140 |
| Spirotetramat | insecticide | 1073000 | 607000 |
| Tebuconazole | fungicide | 200000 | 83050 |
| Tebufenozide | insecticide | 234000 | 100000 |
| Tebuthiuron | herbicide | 30000 | NA |
| Tetraconazole | fungicide | 63000 | 130000 |
| Thiabendazole | fungicide | 34000 | 4000 |
| Thiamethoxam | insecticide | 24 | 5 |
| Thiobencarb | herbicide | 100000 | 100000 |
| Thiophanate methyl | fungicide | 100000 | 114700 |
| Triadimefon | fungicide | NA | 25000 |
| Triadimenol | fungicide | 200000 | 224800 |
| Trichlorfon | insecticide | 59800 | 225 |
| Trifloxystrobin | fungicide | 100000 | 110000 |

Table S4. Studies from Lepidoptera literature review.

| Authors | Year | Species | Compound |
| --- | --- | --- | --- |
| Su et al. | 2014 | *Cnaphalocrocis medinalis* | Azoxystrobin |
| Grafton-Cardwell et al. | 2008 | *Marmara gulosa* | Buprofezin |
| Nasr et al. | 2010 | *Spodoptera littoralis* | Buprofezin |
| Abivardi et al. | 1999 | *Cydia pomonella* | Carbaryl |
| Nagarkatti et al. | 2002 | *Endopiza viteana* | Carbaryl |
| Grafton-Cardwell et al. | 2008 | *Marmara gulosa* | Carbaryl |
| Liu et al. | 2017 | *Agrotis ipsilon* | Chlorantraniliprole |
| Liu et al. | 2017 | *Helicoverpa armigera* | Chlorantraniliprole |
| Pasquini et al. | 2017 | *Lobesia botrana* | Chlorantraniliprole |
| Liu et al. | 2017 | *Spodoptera litura* | Chlorantraniliprole |
| Liu et al. | 2018 | *Bombyx mori* | Chlorantraniliprole |
| Bosch et al. | 2018 | *Cydia pomonella* | Chlorantraniliprole |
| Bird and Walker | 2018 | *Helicoverpa punctigera* | Chlorantraniliprole |
| Lutz et al. | 2018 | *Spodoptera cosmioides* | Chlorantraniliprole |
| He et al. | 2019 | *Agrotis ipsilon* | Chlorantraniliprole |
| Jallow et al. | 2019 | *Tuta absoluta* | Chlorantraniliprole |
| Krishnan et al. | 2020 | *Danaus plexippus* | Chlorantraniliprole |
| Schultz et al. | 2016 | *Euphydryas colon* | Clethodim |
| Schultz et al. | 2016 | *Euphydryas editha* | Clethodim |
| Schultz et al. | 2016 | *Euphydryas phaeton* | Clethodim |
| Pecenka and Lundgren | 2015 | *Danaus plexippus* | Clothianidin |
| Ding et al. | 2018 | *Agrotis ipsilon* | Clothianidin |
| Basley and Goulson | 2018 | *Polyommatus icarus* | Clothianidin |
| Sang et al. | 2016 | *Spodoptera litura* | Cyantraniliprole |
| Xu et al. | 2017 | *Agrotis ipsilon* | Cyantraniliprole |
| Dong et al. | 2017 | *Helicoverpa assulta* | Cyantraniliprole |
| Xu et al. | 2017 | *Ostrinia furnacalis* | Cyantraniliprole |
| Van Laeke et al. | 1991 | *Spodoptera exigua* | Diflubenzuron |
| Grafton-Cardwell et al. | 2008 | *Marmara gulosa* | Diflubenzuron |
| Waldstein et al. | 2000 | *Choristoneura rosaceana* | Fipronil |
| Durham et al. | 2001 | *Ostrinia nubilalis* | Fipronil |
| Gu et al. | 2010 | *Plutella xylostella* | Fipronil |
| Grafton-Cardwell et al. | 2008 | *Marmara gulosa* | Imidacloprid |
| Ahmad et al. | 2013 | *Helicoverpa armigera* | Imidacloprid |
| Krischik et al. | 2015 | *Danaus plexippus* | Imidacloprid |
| Krischik et al. | 2015 | *Vanessa cardui* | Imidacloprid |
| Whitehorn et al. | 2018 | *Pieris rapae* | Imidacloprid |
| James | 2019 | *Danaus plexippus* | Imidacloprid |
| Smagghe et al. | 2003 | *Spodoptera exigua* | Methoxyfenozide |
| Rodriguez Enriquez et al. | 2010 | *Spodoptera exigua* | Methoxyfenozide |
| Zarate et al. | 2011 | *Spodoptera frugiperda* | Methoxyfenozide |
| Saber et al. | 2013 | *Helicoverpa armigera* | Methoxyfenozide |
| Rehan et al. | 2015 | *Spodoptera litura* | Methoxyfenozide |
| Chen et al. | 2019 | *Spodoptera exigua* | Methoxyfenozide |
| Waldstein et al. | 2000 | *Choristoneura rosaceana* | Spinosad |
| Yin et al. | 2008 | *Plutella xylostella* | Spinosad |
| Wang et al. | 2009 | *Helicoverpa armigera* | Spinosad |
| Wang et al. | 2013 | *Spodoptera exigua* | Spinosad |
| Moustafa | 2016 | *Mamestra brassicae* | Spinosad |
| Ahmed | 2016 | *Spodoptera littoralis* | Spinosad |
| Su et al. | 2014 | *Cnaphalocrocis medinalis* | Tebuconazole |
| Waldstein et al. | 2000 | *Choristoneura rosaceana* | Tebufenozide |
| Biddinger et al. | 2006 | *Platynota idaeusalis* | Tebufenozide |
| Fiaz et al. | 2018 | *Anticarsia gemmatalis* | Tebufenozide |
| Yue et al. | 2003 | *Ostrinia nubilalis* | Thiamethoxam |
| Jones et al. | 2012 | *Grapholita molesta* | Thiamethoxam |
| Brown | 1987 | *Trichoplusia ni* | Thiobencarb |

Table S5. Number of exceedances of honeybee LD_50_ concentrations by land use type.

| Land Use Type | Compounds | Exceedance Type | Number of Exceedances |
| --- | --- | --- | --- |
| Agriculture | Clothianidin | Contact | 11 |
| Agriculture | Clothianidin | Oral | 26 |
| Agriculture | Imidacloprid | Oral | 3 |
| Agriculture | Thiamethoxam | Contact | 4 |
| Agriculture | Thiamethoxam | Oral | 13 |
| Production | Thiamethoxam | Contact | 1 |
| Production | Thiamethoxam | Oral | 1 |
| Retail | Cyantraniliprole | Contact | 6 |
| Retail | Cyantraniliprole | Oral | 6 |
| Retail | Thiamethoxam | Contact | 1 |
| Retail | Thiamethoxam | Oral | 3 |
| Roadside | Thiamethoxam | Contact | 1 |
| Roadside | Thiamethoxam | Oral | 1 |
| Urban | Fipronil | Contact | 1 |
| Urban | Fipronil | Oral | 1 |

Figure S1

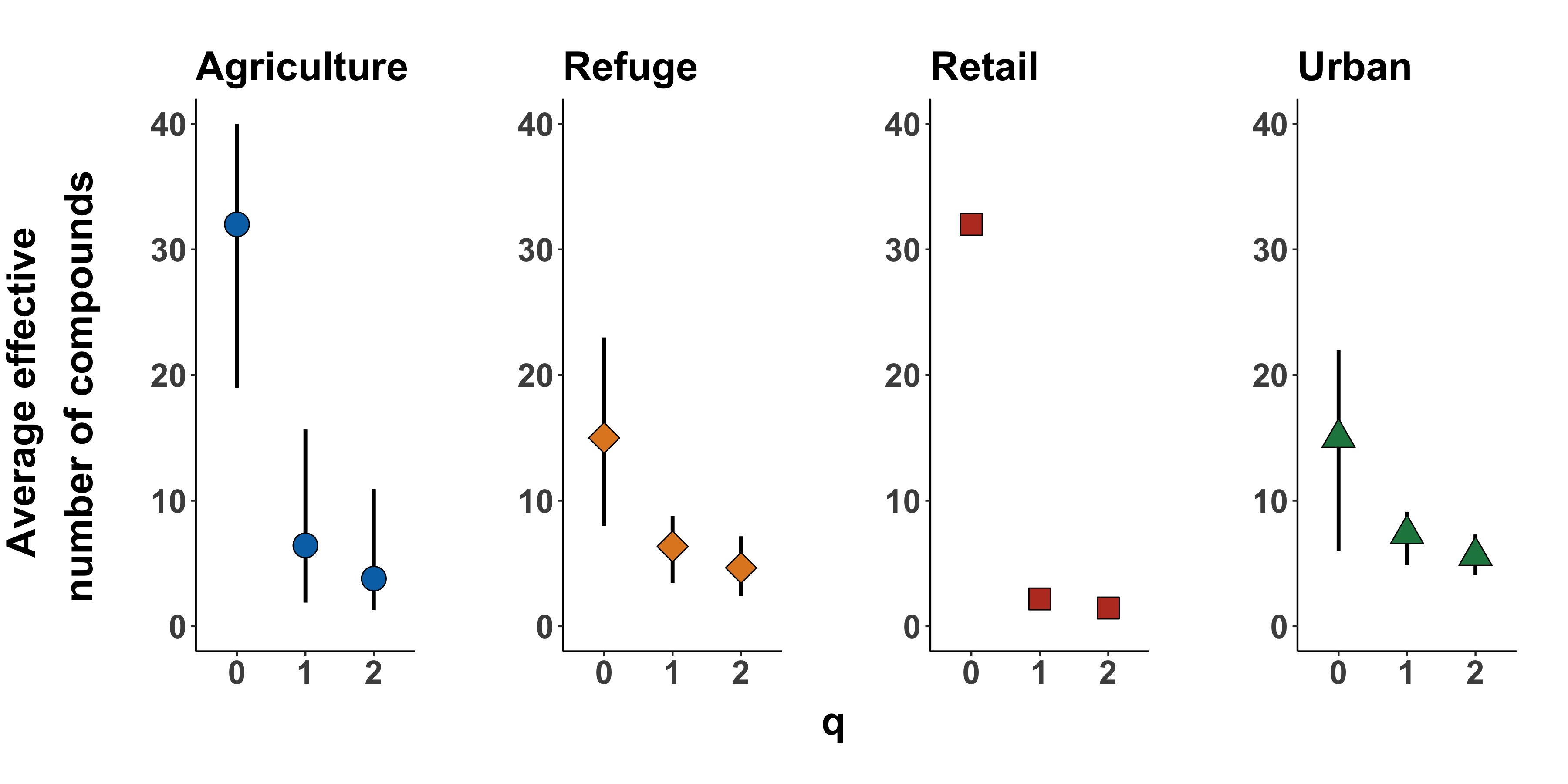

Figure S1. Mean effective numbers of pesticides per sample by land use type using different hill numbers after rarefaction. Points show the mean effective number of compounds per sample. Error bars show the range of effective numbers of pesticides across samples within one land use type.

Figure S2

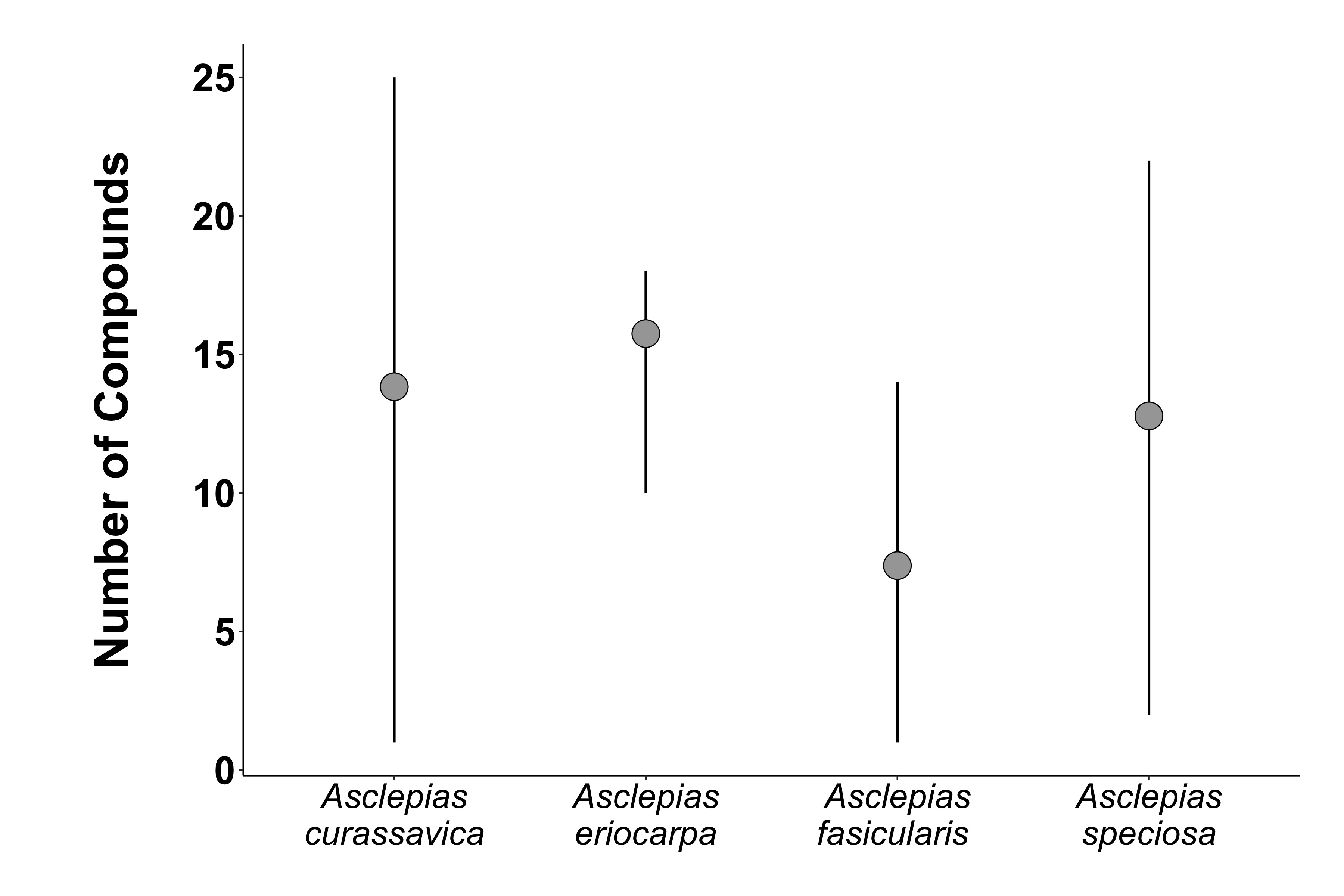

### Figure S2. Variation in the number of compounds per sample by milkweed species. Bars show the maximum and minimum number of compounds detected in any single sample.

Figure S3

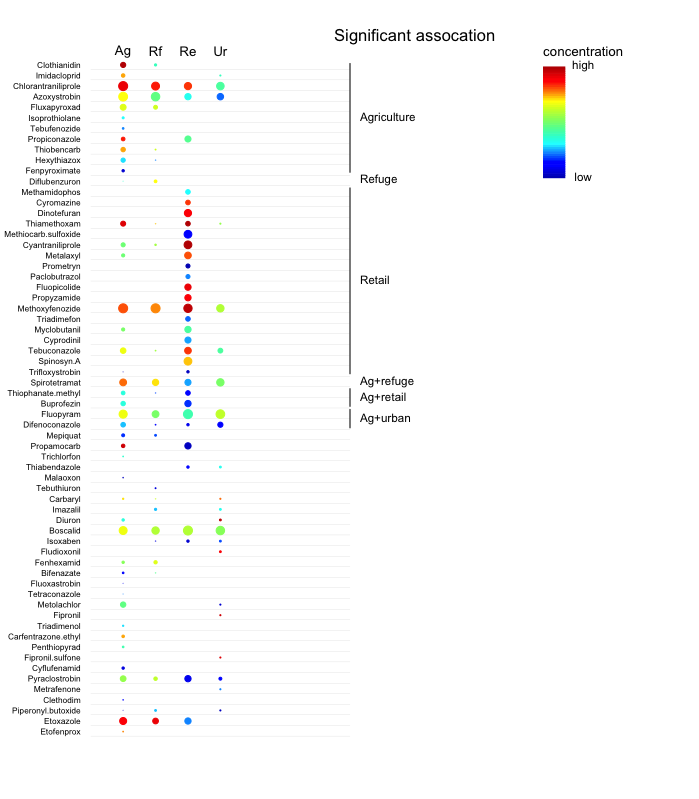

Figure S3. Indicator species analysis examining associations between chemicals and land use types. Color indicates concentration and size the scaled frequency of occurrence. Significant associations are labeled with a black bar and the land use type they are associated with. No correction was made for multiple comparisons.
